# Mitochondrial genetics defines anti-tumour immunity through mitochondrial ROS and PD-1 signalling

**DOI:** 10.64898/2025.12.07.691020

**Authors:** Gennaro Prota, Raquel Justo-Méndez, Uzi Gileadi, Macarena De Andrés-Laguillo, Jose Luis Cabrera-Alarcón, María del Mar Muñoz-Hernández, Valeria Lupi, Sofía C Khouili, Raquel Martínez-De-Mena, Juan Pellico, Rui Benedito, Jesús Ruíz-Cabello, David Sancho, Jose Antonio Enríquez, Ana Victoria Lechuga-Vieco

## Abstract

As central players in cell metabolism, mitochondria influence numerous aspects of health and disease, including the initiation and progression of cancer. Although mitochondrial DNA (mtDNA) mutations have been extensively documented in human cancers for decades, the functional impact of mitochondrial haplogroups on tumour biology remains largely unexplored. Here, we investigate the role of mitochondrial variability in tumour biology using conplastic mouse strains, which are animal models with identical nuclear genomes but different mtDNA haplotypes. We showed that the physiologically relevant variation in mitochondrial ROS (mROS) generation, associated with specific clusters of mtDNA single nucleotide polymorphisms (SNPs), modulated immune responses within the tumour microenvironment and altered tumour growth. We observed strain-dependent differences in the abundance of multiple immune subsets and in PD-1 expression in tumour-infiltrating lymphocytes (TILs). In addition, mtDNA haplotypes influenced cancer progression by modulating tumour angiogenesis through an mROS-independent mechanism. These findings connect nucleo-mitochondrial genetic variability to tumour progression, *de novo* vessel formation and anti-tumour immunity. Tumour immunotherapies should incorporate the spatial and temporal dynamics of cancer evolution and consider mitochondrial genetics as a targetable layer influencing treatment efficacy.

## Main

The multidirectional interaction between tumour transformation, vascular phenotype, and antitumour immunity is a complex biological question that is far from being fully understood. Metabolic changes are correlated with changes in cytokine production and can ultimately influence the ability of infiltrating lymphocytes to find and eliminate cancer cells^1^. In turn, antitumour immunity is directly associated with tumour vascularisation. *De novo* vessels are formed to facilitate nutrient and oxygen delivery in the tumoral area, promoting the proliferation and expansion of cancer cells, while equally promoting improved immune cell infiltration within the tumour, with subsequent arrest of tumour growth^2^. Vascular endothelial growth factor-A (VEGF-A) expression has been associated with increased numbers of tumour-infiltrating lymphocytes (TILs) and programmed cell death ligand 1 (PD-L1) expression in cancer^3^. Elevated reactive oxygen species (ROS) levels, especially in the tumour microenvironment, can drive PD-L1 expression on cancer cells, thereby enhancing immune evasion via the PD-1/PD-L1 interface^4^. Cytotoxic CD8⁺ T cells upregulate PD-1 upon activation, serving as a brake on T cell responses and contributing to T cell exhaustion^5^. Moreover, PD-1 signalling in T cells can further increase intracellular ROS levels, influencing T cell survival and sensitivity to metabolic stress^6^. At the same time, cytoplasmic ROS can activate mitochondrial function in TILs, thereby inhibiting tumour progression through PD-1 blockade^7^. A key objective in advancing anti-tumour therapies is to develop a comprehensive understanding of the spatial and temporal dynamics underlying cancer progression.

Given the key role of mitochondria in metabolic regulation, ROS signalling, and apoptosis^8^, elucidating how mitochondrial DNA (mtDNA) variants shape nuclear gene expression and cellular responses may reveal critical vulnerabilities in tumour biology and immune evasion mechanisms. In humans, mtDNA is a circular double-stranded DNA that encodes 37 genes, including 13 protein components of the mitochondrial electron transport chain (mtETC). Most of the mitochondrial proteome is encoded by the nuclear genome; thus, coordination between the two genomes is required for the correct and functional assembly of the mtETC respiratory complexes. Mutations in mtDNA can lead to diseases such as Leigh syndrome, Leber hereditary optic neuropathy (LHON), mitochondrial encephalomyopathy with lactic acidosis and stroke like episodes (MELAS) and myoclonus epilepsy with ragged red fibres (MERFF)^9^. Beside pathological mutations, mtDNA accumulate single nucleotide polymorphisms (SNPs) that are selected by a discrete maternal lineage during evolution, grouping populations into different mitochondrial haplogroups^10,11^. Determining the potential functional relevance of mtDNA SNPs is complex. It is well known that mtDNA influences numerous aspects of pathophysiology, including the initiation and progression of cancer^12–15^. While a substantial amount of data correlating mtDNA mutations with cancer is available from sequencing studies, only a few functional studies have investigated the potential functional role of mitochondrial haplogroups and haplotypes in cancer, representing a crucial knowledge gap^16–21^. Studies employing transmitochondrial cybrids^22,23^ or MNX mice^24,25^,have investigated the role of mitochondrial haplogroups on the metastatic potential of tumour cells in mice in the context of metastasis, identifying a prominent role of ROS.

The association between mtDNA variants in cancer cells and the efficacy of immunotherapy has only recently been described^26,27^. Mutations in cancer cell-derived mtDNA can reshape the tumour microenvironment, thereby altering the immune landscape and influencing tumour sensitivity to PD-1 checkpoint blockade^27^. This raises the question of how the presence of mitochondrial SNPs within immune cells affects tumour-immune interactions. While the role of mitochondria in orchestrating both innate and adaptive immune responses is well established^28^, our understanding of how physiological mitochondrial variants modulate immune cell function is limited. To date, only a few studies have reported associations between specific mtDNA haplogroups and disease susceptibility or immune phenotypes. Human haplogroup H is associated with better outcomes after severe sepsis^29^ and better sperm motility^30^, and has also been linked to protection against SARS-CoV2 severity^31^ and enhanced recovery after antiviral drug treatment in HIV-infected patients^32^. Furthermore, mitochondrial haplogroup A has been proposed as a risk factor for acquired immunodeficiency syndrome (AIDS)^33^ and haplogroup D5 for chronic hepatitis B virus (HBV)^34^. Mitochondrial ROS (mROS) also play important roles as secondary messengers in T-cell activation and defence against cancer^35,36^ with several studies suggesting that they increase angiogenesis by upregulating VEGF expression in endothelial cells^37^. Although differences in basal mROS generation and handling have been reported in both cells^38–40^ and murine models^41–43^ carrying maternally inherited mitochondrial polymorphisms on an identical nuclear background, the impact of mitochondrial genetic variation on immune cell biology is only beginning to be explored^26,44,45^. Mitochondrial haplotypes in mice can impair regulatory T cell (Treg) function through increased ROS generation, enhancing Treg activity, leading to enhanced tumour rejection^45^. Moreover, HG-T human mitochondrial haplogroups are predictive of resistance to immune checkpoint therapies in melanoma, linked to altered ROS detoxification pathways, and distinct baseline CD8^+^ T cell phenotypes^26^. The specific role of mROS in driving tumour progression through effects on the vascular phenotype and antitumour immunity, and the mechanisms by which mtDNA variants shape these processes remain undefined.

Here, we demonstrate that mitochondrial SNPs shape the antitumour immune response by modulating angiogenesis, immune cell infiltration, and T cell activation. Using conplastic mouse strains, we identified a novel mechanistic role for mROS in tumour progression and responsiveness to immune checkpoint blockade. Importantly, genetically targeted modulation of mROS in T cells altered tumour progression and PD-1 signalling, identifying a direct link between mitochondrial genetics, immune regulation, and therapeutic outcomes. These findings highlight mitochondrial haplotypes as determinants of tumour-immune interactions and suggest that stratifying patients by mtDNA background could inform personalised approaches to checkpoint immunotherapy.

## Results

### Enhanced tumour development and progression of fibrosarcoma and melanoma in BL/6^C57^ compared to BL/6^NZB^ conplastic mice

Tumour progression depends on mtDNA haplotypes in mice, with distinct mitochondrial origins differentially shaping tumour growth and immune responses^45^. C57BL/6 mice demonstrate an age-dependent increase in spontaneous tumour incidence that follows an exponential trajectory^46,47^. We have previously reported a tendency to develop a lower frequency of tumours in animals with identical nuclear background (C57BL/6), but with NZB-derived instead of C57 mtDNA^42^. Thus, we first confirmed a higher incidence of spontaneous tumours in mice that carry C57 mtDNA, analysed in experimental groups with 80-90 weeks of age (Fig. 1a). This could reflect either an mtDNA-associated predisposition to tumour formation or mtDNA-associated differences in tumour progression. In humans, it has been shown an association between mtDNA haplogroup and the response to immune checkpoint inhibitors in advanced-stage melanoma^26^. To investigate whether mtDNA variation also influences tumour predisposition in humans, we analysed the prevalence of mitochondrial haplogroups in patient cohorts. Using UK Biobank data^48^ as a population reference, we compared macro-haplogroup frequencies with those observed in patient-derived samples from the CheckMate-067 trial (NCT01844505)^26^. For this purpose, a binomial test was performed for each macro-haplogroup considered in this study: H, U, K, T, J, as well as a composite group of other European haplogroups, which includes W, V, R, I, and X, as shown in Extended Data Fig. 1a and detailed in Supplementary Table 1. All considered European macro-haplogroups were found to be distributed as expected, showing no statistically significant bias when compared to the reference UK population, after correction for multiple testing (corrected p < 0.05) using the Benjamini-Hochberg method. These results indicate that the mtDNA macro-haplogroup distribution in the CheckMate-067 cohort is representative of the broader UK-European genetic background, suggesting minimal population stratification bias and providing no evidence for enrichment of specific mtDNA haplogroups consistent with melanoma predisposition.

**Figure 1.**
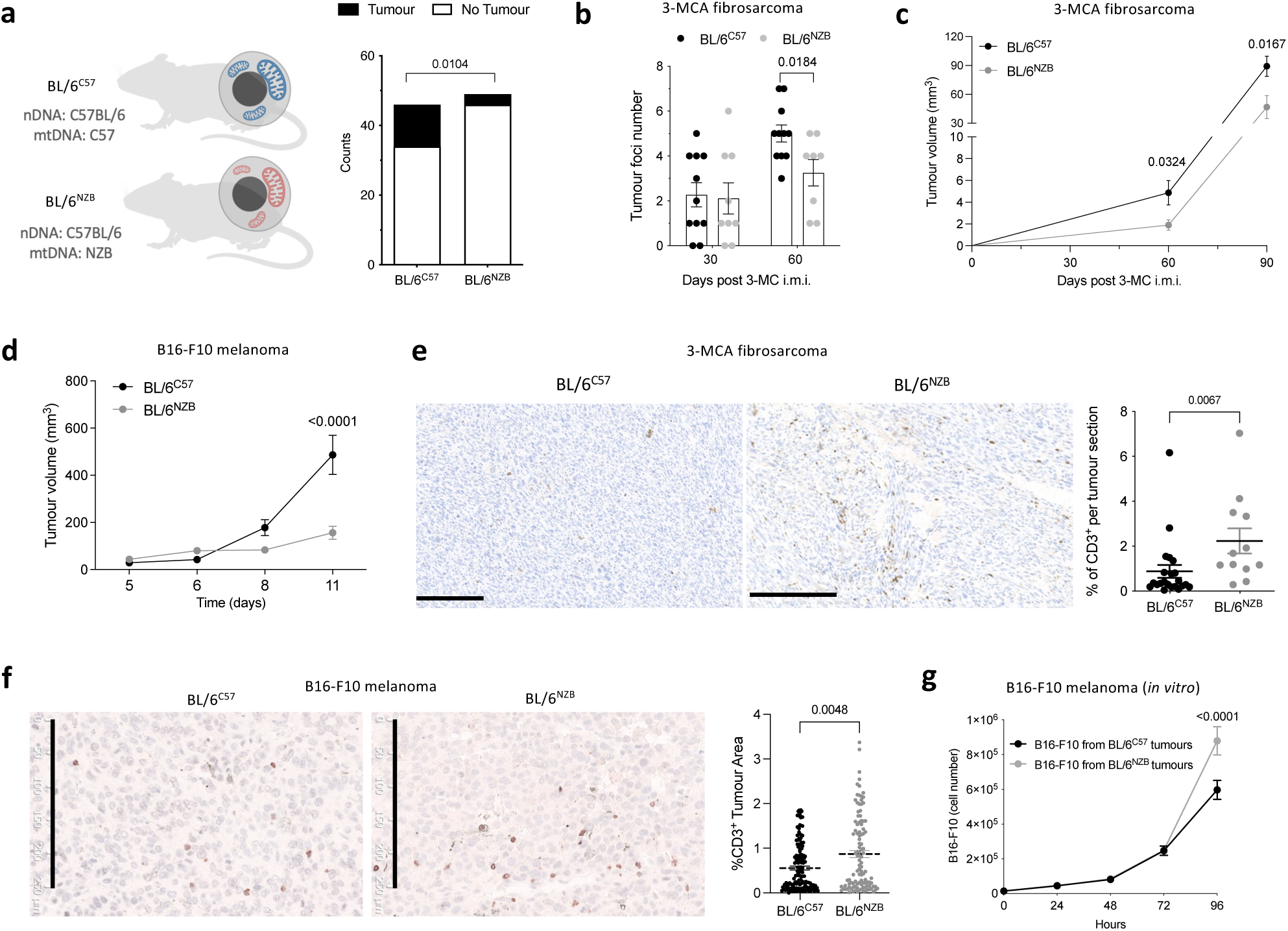
Age-related spontaneous tumour incidence and characterisation of chemically induced fibrosarcoma and B16-F10 melanoma in conplastic strains. **a,** Schematic representation of BL/6^C57^ and BL/6^NZB^ mice (left panel) and spontaneous hepatic tumour incidence in 80-100 weeks old mice (n= 46 BL/6^C57^ and 49 BL/6^NZB^ mice) (right panel). Fisher’s exact test analysis. **b-c,** Mice received intramuscular injections of 3-MCA and the number of tumours per rear leg (b) and tumour progression (c) were shown at the indicated time points (n= 8-12 mice per genotype). Two-tailed unpaired t test. **d,** Tumour growth measurements of the two conplastic strains after received subcutaneous injections with B16-F10 cells (n= 20-24 mice per genotype; four independent experiments). Two-way ANOVA with Šídák’s multiple comparisons test. **e,** Representative images of immunohistochemical sections of fibrosarcoma tumours collected on day 90 after 3-MCA injection of BL/6^C57^ and BL/6^NZB^ mice, stained for CD3 (left panel), and quantified based on the percentage of CD3^+^ area (right panel), with each dot representing a different tumour section (n= 5-6 mice per genotype). Scale bar: 250μm. Kolmogorov-Smirnov nonparametric test. **f,** Representative images of immunohistochemical sections of melanoma tumours from BL/6^C57^ and BL/6^NZB^, stained for CD3 (left panel), and quantified based on the percentage of CD3^+^ area (right panel). Each dot represent*s* a different tumour section (n= 15 BL/6^C57^ mice and n= 11 BL/6^NZB^ mice; two independent experiments). Kolmogorov-Smirnov nonparametric test. Scale bar: 250μm. **g,** *In vitro* proliferation analysis of B16-F10 cells isolated from tumours, taken at day 14 after subcutaneous injection (n= 12 tumours from BL/6^C57^ and n= 10 tumours from BL/6^NZB^ mice). Two-way ANOVA with Šídák’s multiple comparisons test. Each dot represents an animal unless otherwise noted. Data are given as means ± SEM. *p-value < 0.05, **p-value < 0.01, and ****p-value < 0.0001.

To test the impact of mtDNA haplotype on tumour development and progression in mice, conplastic animals were injected intramuscularly with 3-methylcholanthrene (3-MCA) to induce fibrosarcoma (Extended Data Fig. 1b), a malignant mesenchymal tumour. Notably, solid tumour formation in this model originates from only one or a few initiating cells. These tumours are driven by activation of K-Ras alone or in combination with N-Ras proto-oncogenes^49,50^ and are highly sensitive to p53 activity, with sustained activation of DNA damage response pathways^51^. We monitored tumour development using magnetic resonance imaging (MRI) in combination with the paramagnetic gadolinium-based contrast agent gadopentetate dimeglumine (Gd-DTPA tradename Magnevist^TM^), which enhances contrast in the tumoral area (Fig. 1b-c and Extended Data Fig. 1c). Initially, we observed an increased frequency of tumoral foci after 60 days of 3-MCA injection in C57 mtDNA mice (Fig. 1b and Extended Data Fig. 1c) and this correlated with accelerated tumour progression (Fig. 1c).

Chemical induction of tumours in conplastic strains do not provide enough information to discern between intrinsic cancer cell tumorigenicity and the ability controlling the tumour progression in conplastic animals. To separate these contributing factors, we subcutaneously injected mice with B16-F10 melanoma cells (Extended Data Fig. 1d). In this orthotopic melanoma model, mtDNA harboured by tumour cells is the same in both BL/6^C57^ and BL/6^NZB^ recipient animals. In line with results observed in fibrosarcoma, the tumoral growth rate is slower in BL/6^NZB^ host mice when compared to that of their BL/6^C57^ counterparts (Fig.1d). Histological staining for Ki-67, a marker of dividing cells, confirmed a lower proportion of Ki67^+^ cells in tumours from BL/6^NZB^ mice (Extended Data Fig. 1e). Because mitochondrial proteins were recently shown to be highly immunogenic^52^, we next investigated whether the reduced growth of B16-F10 tumours in BL/6^NZB^ recipient mice could result from immune rejection. Specifically, we considered whether differences in mtDNA between BL/6^NZB^ hosts and the B16-F10 cells, originally derived from C57BL/6 mice^53^, might contribute to this effect. For that, we generated B16-F10 transmitochondrial cybrid cells (B16-F10^NZB^) harbouring the original B16-F10 nuclear DNA (nDNA) but with NZB mtDNA and performed subcutaneously injections in BL/6^NZB^ and BL/6^C57^ mice (Extended Data Fig. 1f). These transmitochondrial cybrid melanoma cells also generated tumours at a lower rate in BL/6^NZB^ mice, reproducing the behaviour of the original B16-F10 melanoma cells (Fig. 1d and Extended Data Fig. 1f). This observation is consistent with the BL/6^NZB^ host background being the primary driver of tumour control, rather than tumour-intrinsic features arising from distinct mitochondrial genetics.

To further dissect the physiological bases of the differences in tumour progression we first focused on the spatial distribution of cells within the tumoral area. Chemical induction of fibrosarcoma is a tumour model used to investigate immune surveillance^54–56^, in which NK and CD8^+^ T cells are controlling the tumour formation and progression^57–59^. We analysed the changes in the level of TILs in both strains 90 days after the 3-MCA tumour induction. We did not observe significant differences in total immune cell infiltration (CD45⁺) between the two strains (Extended Data Fig. 1g), although animals carrying NZB mtDNA had higher number of TILs (total CD3⁺) than BL/6^C57^ mice (Fig. 1e). However, mechanisms that control chemical carcinogenesis are different from the transplanted tumour cells^60^. Thus, we tested the differences in CD3⁺ T cell infiltration in our B16-F10 melanoma model (Fig. 1f). Interestingly, we observed the same patterns observed with the 3-MCA model. The lower tumour volume in conplastic NZB mtDNA mice correlated with a higher T cell infiltration within the melanoma tumours when compared to wild-type C57BL/6 mice (Fig. 1f). After re-isolating melanoma cells from excised BL/6^C57^ and BL/6^NZB^ tumours and culturing them *in vitro*, we found that the *in vivo* tumour growth differences were abolished (Fig. 1g). Opposite to the *in vivo* behaviour, B16-F10 tumours excised from mice harbouring C57 mtDNA had a slower proliferation rate compared to tumour cells excised from BL/6^NZB^ mice *in vitro* (Fig. 1g), pointing out the role of the tumour microenvironment in the *in vivo* differential tumour growth.

Altogether, our data demonstrate that the mtDNA haplotype of recipient mice significantly influenced tumour development and progression dynamics. Both the chemically induced fibrosarcoma model and the orthotopic melanoma model showed that that NZB versus C57 mtDNA backgrounds were associated with different tumour growth rates, accompanied by differences in intratumoral T-cell infiltration.

### The match between nuclear and mitochondrial DNA influences T cell phenotypes within the tumour microenvironment

Our first results obtained are in line with the notion that TILs are important to control tumour growth and mitochondrial genetics might affect the immune cytolytic activity. To understand how the anti-tumoral immunity is shaped in mice with different mtDNA haplotypes, we performed analysis using Fluorescence-activated Cell Sorting (FACS) to explore possible differences in CD8^+^ and CD4^+^ T cell subsets and the expression of the programmed cell death protein 1 (PD-1) marker in both, 3-MCA-induced fibrosarcoma (Fig. 2a and Extended Data Fig. 2a and c) and B16-F10 melanoma (Fig. 2b and Extended Data Fig. 2b and d) tumour models. The quantitative characterisation made by FACS showed a higher frequency of CD8^+^ T cells infiltrating both type of tumours in BL/6^NZB^ compared to BL/6^C57^ (Fig. 2a-b, left panels) while no differences were observed in the CD4^+^ T cell population between the two strains (Extended Data Fig. 2c and d). These results suggests that CD8^+^ T cells are important mediators of the anti-tumour immune response in our conplastic models. Furthermore, we observed an increase of effector CD8^+^CD44^+^PD-1^+^ cells in BL/6^NZB^ mice in both tumour models (Fig. 2a and b, right panels). The expression of PD-1 is linked to an exhaustion status of T cells in the latest stage of tumour progression^61^, but its detection at an early/intermediate stage is a proof of immune activation^62^. Consistently with that result, we observed upregulation of PD-L1 on B16-F10 melanoma cells in BL/6^NZB^ conplastic mice compared to BL/6^C57^ that could be mediated by the presence of IFN-γ and TNF-α within the tumour microenvironment (Extended Data Fig. 2e-f).

**Figure 2.**
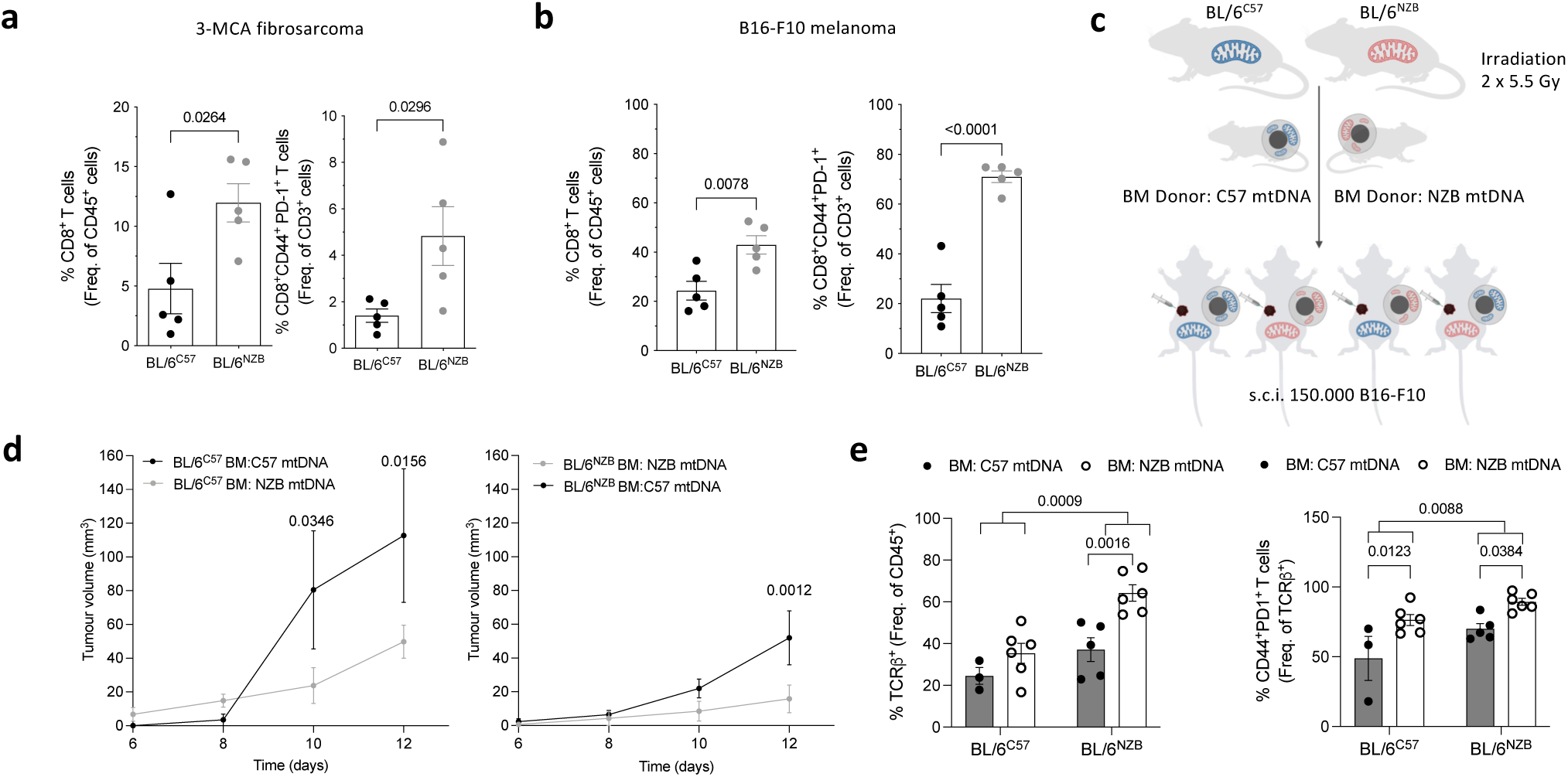
Characterisation of antitumour responses in conplastic mouse strains. **a-b,** FACS analyses of tumour-infiltrating lymphocytes (TILs) in 3-MCA-induced-fibrosarcomas (**a**) or B16-F10 melanoma tumours (**b**), showing the percentage of CD8^+^ T cells within total CD45^+^ immune cells (left panel), and the percentage of effector CD8^+^ T cells expressing PD-1 within TILs (right panel) (n= 5 mice per genotype). Unpaired t test. Data are representative of 3 independent experiments. **c,** Schematic representation of the bone marrow (BM) transplantion experimental design. Groups of BL/6^C57^ and BL/6^NZB^ mice were irradiated, and each strain was repopulated with the BM from BL/6^C57^ or BL/6^NZB^ mice covering all the possible combinations. 8-10 weeks after BM transplant, mice were subcutaneously injected (s.c.i) with B16-F10 melanoma cells. **d,** B16-F10 melanoma tumour growth measured at the indicated time points (n=5-6 mice per experimental group). Two-way ANOVA with Šídák’s multiple comparisons test. Data are representative of 3 independent experiments. **e,** FACS analysis in TILs showing the percentage of TCRβ^+^ cells within total CD45^+^ immune cells (left panel), and the percentage of effector CD8^+^ T cells expressing PD-1 within total TILs (right panel) (n=3-6 mice per group). Two-way ANOVA with Šídák’s multiple comparisons test. Data are representative of 3 independent experiments. Each dot represents an animal unless otherwise noted. Data are given as means ± SEM. *p-value < 0.05, **p-value < 0.01, ***p-value < 0.001, and ****p-value < 0.0001.

We then sought to determine if the nDNA/mtDNA interaction was mediating the control of tumour growth acting on immune cells or if there were other non-immune cell mediated factors. To this aim we irradiated BL/6^C57^ and BL/6^NZB^ mice and replaced their bone marrow (BM) covering all the possible combinations (Fig. 2c). 8-10 weeks post BM transplantation, we challenged the mice with B16-F10 melanoma cells and monitored the tumour growth (Fig. 2d). This approach led us to dissect the role of the different mtDNA in the immune compartment. Results showed that host mice with BM from BL/6^NZB^ conplastic mice had a superior ability to control tumour growth compared with mice that received the BM from BL/6^C57^ mice (Fig. 2d). This result pointed out the role of mtDNA/nDNA interaction in the generation of an effective immune response. TILs analyses showed higher ability of T cells from BL/6^NZB^ to infiltrate tumours (Fig. 2e, left panel), and higher expression of PD-1 on NZB mtDNA effector T cells compared to the BL/6^C57^ strain (Fig. 2e, right panel), in line with a more aggressive immune response.

These findings support the role of TILs in controlling tumour progression and highlight mitochondrial genetics as a determinant of immune cytotoxic activity. BL/6^NZB^ mice showed enhanced CD8^+^ T cell infiltration, higher PD-1 expression, and improved tumour control. Bone marrow transplantion experiments confirmed that these effects are immune-mediated and driven by mtDNA/nDNA interactions in the haematopoietic compartment.

### T-cell mitochondrial ROS suppress tumour progression through PD-1 regulation

We previously described that conplastic mouse models present different capacity of ROS handling with an increase in the basal mROS levels in mice harbouring NZB mtDNA^42^. Moreover, the overexpression of MnSOD (manganese superoxide dismutase) has been reported to promote tumorigenicity^63^ and enhance the metastatic properties of the cancer cells in different types of tumours^64,65^. However, the contribution of host-derived mROS, as opposed to tumour-intrinsic mROS to cancer progression, has not been previously defined.

We quantified total intracellular ROS levels in TILs from BL/6^NZB^ and BL/6^C57^ tumours and confirmed increased ROS in cells harbouring NZB mtDNA (Fig. 3a, and Extended Data Fig. 3b). To assess the functional relevance of ROS in tumour control, we injected BL/6^C57^ mice subcutaneously with B16-F10 cells and treated them with either a global ROS scavenger (N-acetylcysteine, NAC) or a mitochondria-targeted ROS scavenger (mitoquinone mesylate, MitoQ) (Fig. 3b-c). Notably, only MitoQ administration altered tumour progression, resulting in increased tumour growth (Fig. 3b). Neither total ROS nor mROS scavenging affected the frequency of intratumoral CD8⁺ or CD4⁺ T cells (Fig. 3c, left panel, and Extended Data Fig. 3a). However, the inhibition of mROS specifically reduced CD8⁺ T-cell activation, as reflected by decreased PD-1 expression (Fig. 3c, right panel).

**Figure 3.**
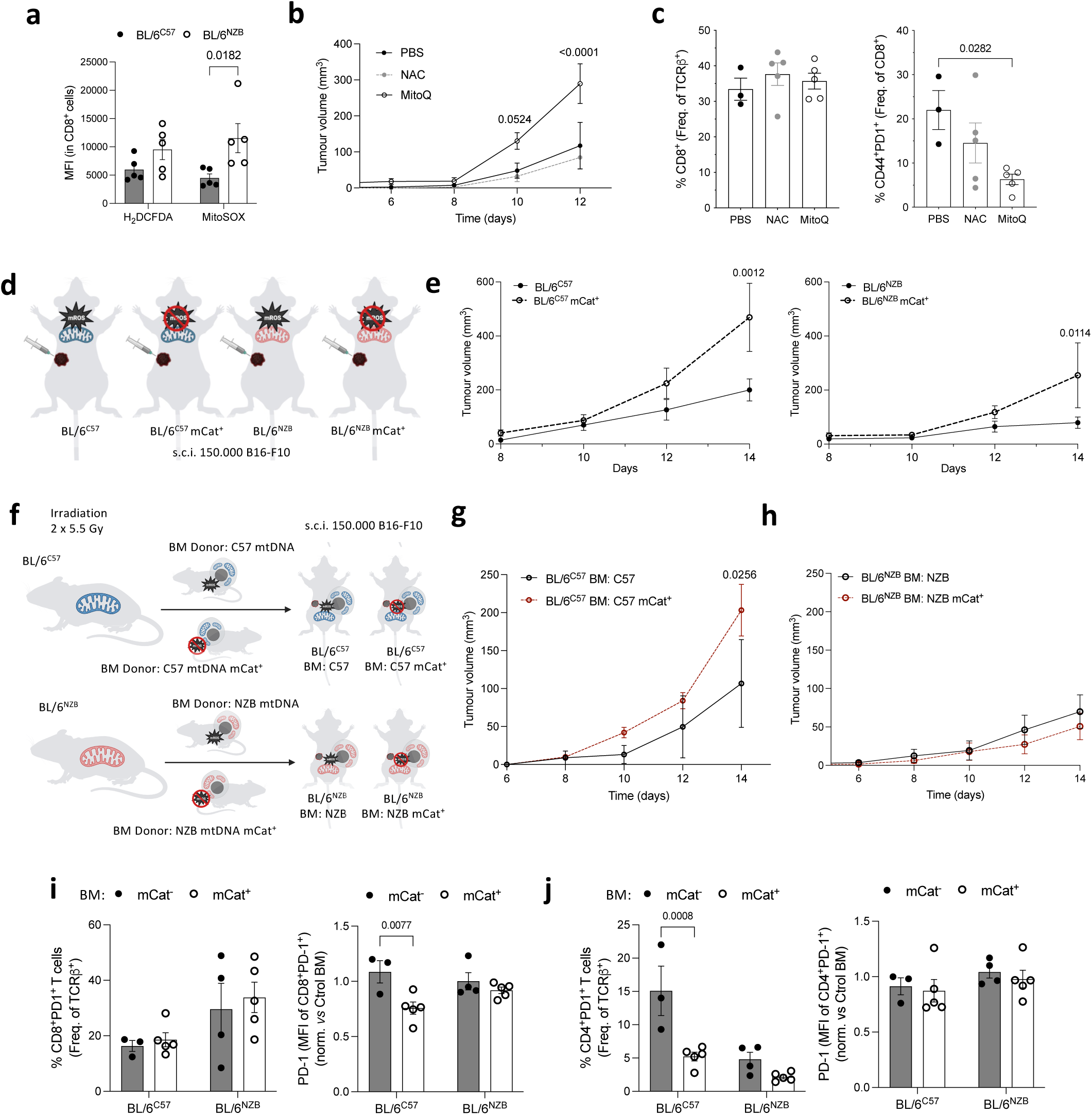
Role of mROS in TILs during tumour progression. a, FACS analysis of total cellular ROS levels (H_2_DCFDA staining) and mROS levels (MitoSOX staining) in CD8^+^ T cells 14 days post injection of B16-F10 melanoma cells in BL/6^C57^ and BL/6^NZB^ mice (n=5 mice per genotype). Two-way ANOVA with Šídák’s multiple comparisons test. **b,** Tumour growth in wild-type C57BL/6 mice injected with B16-F10 melanoma cells and the indicated treatments (n= 5 NAC, N-acetyl cysteine; n= 5 MitoQ; n= 3 PBS - vehicle). Two-way ANOVA with Dunnett’s multiple comparisons test. **c,** FACS analysis of TILs showing the frequency of CD8^+^ T cells (left panel) and the percentage of effector CD8^+^ T cells expressing PD-1 (right panel). Each dot represents an individual tumour from the indicated treatment groups (n= 5 NAC, N-acetyl cysteine; n= 5 MitoQ; n= 3 PBS - vehicle). One-way ANOVA with Dunnett’s multiple comparisons test. **d,** Schematic representation of the orthotopic B16-F10 melanoma model in conplastic mice with or without expression of the mitochondrial-target catalase (mCat^+^). **e**, Tumour growth measured at the indicated time points for the specified genotypes (n=6-8 mice per group). Two-way ANOVA with Šídák’s multiple comparisons test. **f,** Schematic of the bone marrow (BM) transplantation design. Conplastic mice were irradiated and reconstituted with bone marrow from BL/6^C57^ or BL/6^NZB^ donors, with or without mitochondria-targeted catalase (mCat⁺). After 8-10 weeks post-BM transplant, mice were subcutaneously injected (s.c.i.) with B16-F10 melanoma cells. **g**, Tumour growth measured at the indicated time points for the specified genotypes (n = 5 mice per experimental group). Two-way ANOVA with Šídák’s multiple comparisons test. **h**, Tumour growth measured at the indicated time points for the specified genotypes (n = 12 control mice and n= 17 mCat^+^ mice per group). Two-way ANOVA with Šídák’s multiple comparisons test. **i**, FACS analysis showing the percentage of CD8^+^ T cells expressing PD-1 in TILs (left panel) and PD-1 expression (MFI) within this subset (right panel) (n=3-5 mice per experimental group). Two-way ANOVA with Šídák’s multiple comparisons test. **j**, FACS analysis showing the percentage of CD4^+^ T cells expressing PD-1 in TILs (left panel) and PD-1 expression (MFI) within this subset (right panel) (n=3-5 mice per experimental group). Two-way ANOVA with Šídák’s multiple comparisons test. Each dot represents an animal unless otherwise noted. Data are given as means ± SEM. *p-value < 0.05, **p-value < 0.01, and ****p-value < 0.0001.

NAC and MitoQ have complex, pleiotropic antioxidant mechanisms and are not selective for a single ROS species. To further dissect the role of mROS, we generated constitutive conplastic mouse models expressing a mitochondria-targeted catalase (mCat)^66^, enabling selective depletion of mitochondrial H_2_O_2_ in the mitochondrial matrix (Fig. 3d). We first evaluated the impact of mCat expression on B16-F10 tumour development and progression. Reduction of mitochondrial H_2_O_2_ in mCat⁺ mice resulted in accelerated tumour growth relative to their corresponding BL/6^C57^ and BL/6^NZB^ controls (Fig. 3e). In parallel, we analysed the TIL landscape as previously described in Fig. 2. The overall abundance of total CD3^+^ T cells was comparable across strains, as shown by immunohistochemical analysis (Extended Data Fig. 3c). However, the proportion of effector T cells expressing PD-1 was significantly reduced only in tumours from BL/6^NZB^ mCat^+^ mice (Extended Data Fig. 3d). By contrast, PD-L1 expression on cancer cells remained unchanged between strains (Extended Data Fig. 3e).

To better understand whether differences in mROS levels within immune cells were major contributors to the observed changes in tumour progression, we carried out BM transplant experiments. Specifically, we generated BM chimaeras in which the tumour-injected cells were identical across groups (B16-F10), ensuring that any differences arose from the host non-haematopoietic compartment or from donor-derived haematopoietic cells (Fig. 3f), including T cells. Lethally irradiated BL/6^NZB^ and BL/6^C57^ mice were reconstituted with BM from either BL/6^C57^, BL/6^NZB^, BL/6^C57^ mCat^+^ or BL/6^NZB^ mCat^+^ donors. Notably, melanoma progression increased significantly in BL/6^C57^ hosts receiving BM from BL/6^C57^ mCat^+^ donors (Fig. 3g), whereas no such effect was observed in BL/6^NZB^ hosts transplanted with BL/6^NZB^ mCat^+^ BM (Fig. 3h). Although the frequency of intratumoral CD8^+^PD-1^+^ cells was unchanged across groups (Fig. 3i, left panel), mCat expression in the haematopoietic compartment of BL/6^C57^ mice reduced PD-1 expression levels (Fig. 3i, right panel). Consistent with this, the same group also showed a lower frequency of PD-1-expressing CD4^+^ T cells (Fig. 3j), suggesting broader attenuation of PD-1-associated activation downstream of mROS signalling. In contrast, BL/6^NZB^ mice showed no detectable changes in TIL phenotypic parameters upon haematopoietic mCat expression (Fig. 3h-j).

Overexpression of a mitochondria-targeted catalase reduces mROS levels in immune cells, leading to diminished PD-1 expression and impaired T-cell effector function. Consistent with this, suppressing mitochondrial ROS signalling in the haematopoietic compartment accelerated B16-F10 melanoma development and progression specifically in BL/6^C57^ mice, which exhibit lower basal mROS levels.

### Mitochondrial DNA haplotype influences tumour angiogenesis independently of mROS signalling

An important factor for tumour development and progression is the ability to generate neo-vasculature^67^. Because host BL/6^NZB^ mice displayed enhanced tumour control compared with BL/6^C57^ mice, regardless of BM donor source (Fig. 2d and Extended Data Fig. 4a), we investigated whether differences in nDNA/mtDNA crosstalk could modulate angiogenesis as a non-immune mechanism contributing to tumour growth. First, we analysed the angiogenic capacity under basal conditions using the two conplastic strains. Using a directed *in vitro* angiogenesis assay (DIVAA), we identified that BL/6^NZB^ mice have a greater ability to generate *de novo* vasculature compared to BL/6^C57^ mice (Fig. 4a).

**Figure 4.**
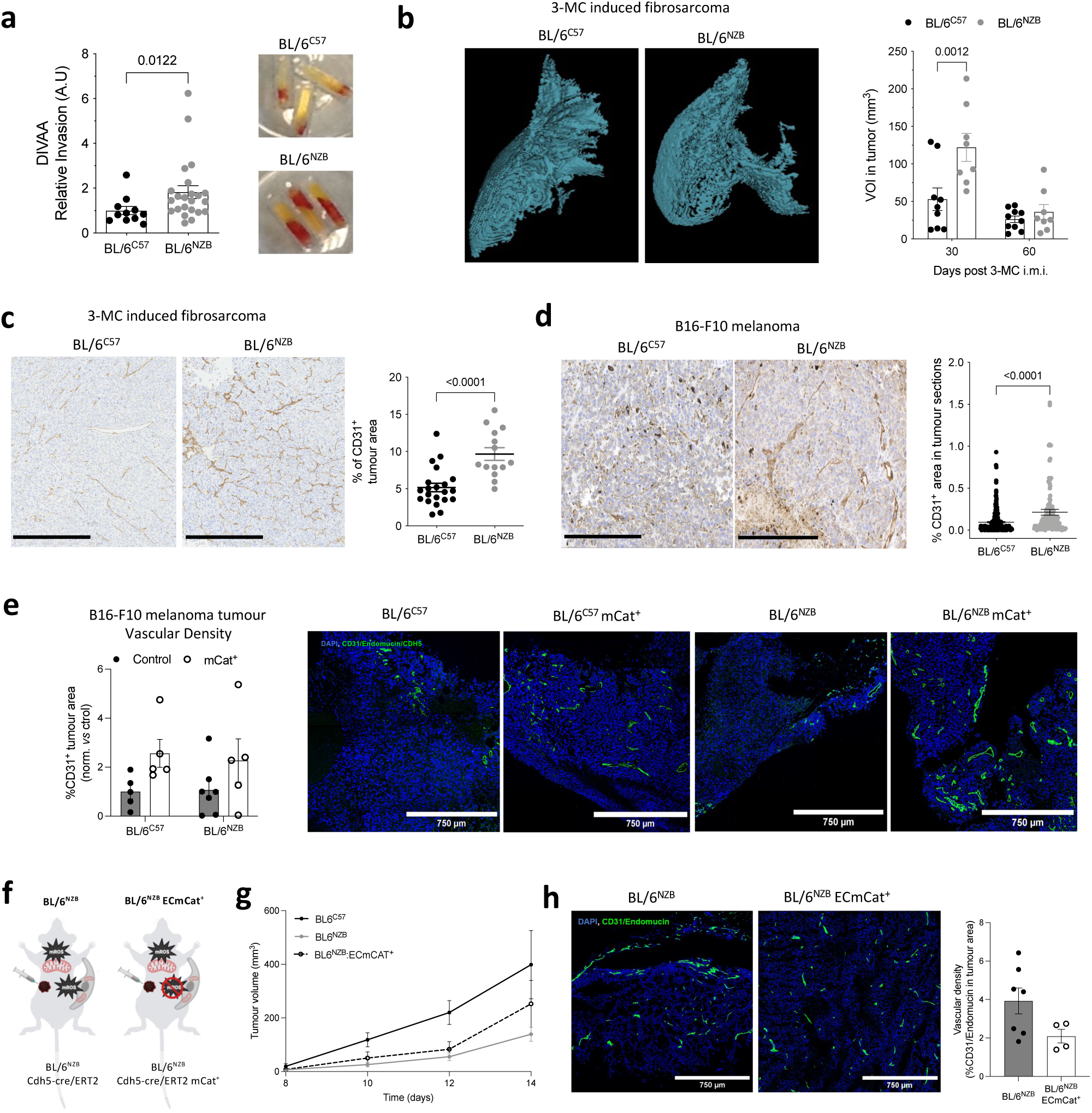
Characterisation of vasculature in tumour bearing mice. **a,** Relative cellular invasion in the directed *in vivo* angiogenesis assay (DIVAA), quantified by fluorescence (A.U.) 20 minutes after intravenous FITC-dextran injection (left panel) and, representative images of recovered angioreactors from both strains (right panel), collected 8 weeks after implantation. Each dot from left panel represents a distinct angioreactor from an individual mouse (n= 11 BL/6^C57^ and n= 23 BL/6^NZB^ mice; two independent experiments). Two-tailed unpaired t test. **b,** 3D MRI reconstructions of fibrosarcomas 30 days after intramuscular 3-MCA injection in BL/6^C57^ and BL/6^NZB^ mice (left panel) and, quantification of total volume of interest at 30- and 60-days post-3-MCA injection (right panel) (n= 8-10 mice per group). Two-way ANOVA with Šídák’s multiple comparisons test. **c,** Representative immunohistochemical images of fibrosarcoma tumour sections collected on day 90 after 3-MCA injection in BL/6^C57^ and BL/6^NZB^ mice, stained for CD31 (left panel), and quantified based on the percentage of CD31^+^ area (right panel), with each dot representing a different tumoral section (n= 5-6 mice per genotype). Two-tailed unpaired t test. Scale bar: 500μm. **d,** Representative images of immunohistochemical images of melanoma tumour sections from BL/6^C57^ and BL/6^NZB^, stained for CD31 (left), and quantified based on the percentage of CD31^+^ area (right), with each dot representing a different tumoral section (n= 15 BL/6^C57^ mice and n= 11 BL/6^NZB^ mice; two independent experiments). Kolmogorov-Smirnov nonparametric test. Scale bar: 250μm. **e,** Quantification of vascular density by immunohistochemistry showing the percentage of CD31^+^ cells per tumour area (left panel), and representative images of vascular density (in green: CD31/Endomucin/CDH5) and nuclei (in blue: DAPI) (right panel) in conplastic mice with or without mitochondria-targeted catalase expression (n= 5-7 mice per group). Two-way ANOVA with Šídák’s multiple comparisons test. Scale bar: 750μm. **f,** Schematic representation orthotopic B16-F10 melanoma model in a newly generated BL/6^NZB^ strain expressing mCat under the promoter Cdh5-cre-ERT2 to selectively reduce mROS only in endothelial cells (ECs). **g,** Tumour growth measurements comparing BL/6^C57^ (n=5), BL/6^NZB^ (n=5), and BL/6^NZB^ expressing mitochondria-targeted catalase (mCat^+^) in tumour endothelial cells (TECs; n=9). Two-way ANOVA with Šídák’s multiple comparisons test. **h,** Representative images of immunohistochemistry showing vascular density (CD31/Endomucin, green) and nuclei stained with DAPI (blue) in BL/6^NZB^ and BL/6^NZB^ ECmCat^+^ mice (left panel) and, the percentage of CD31^+^ area in tumour sections (right panel) (n= 5-7 mice per group). Two-tailed unpaired. Scale bar: 750μm. Each dot represents an animal unless otherwise noted. Data are given as means ±SEM. *p-value < 0.05, **p-value < 0.01, and ****p-value < 0.0001.

To explore whether this difference in angiogenesis profile can be applied to tumour biology, we next analysed the vascular pattern of 3-MCA fibrosarcoma tumours using MRI, comparing between the conplastic strains (Fig. 4b). MRI using gadolinium-based contrast agents is an effective method of monitoring tumour responses based on both morphological and vascular patterns^68^. Using this technique, which provides positive contrast enhancement in intravascular and interstitial areas, we visualised clear differences between the mouse strains in the density and spatial distribution of fibrosarcoma peritumoral vessels (Fig. 4b, left panel). Quantification of total volume of interest (VOI) showed increased vessel density in BL/6^NZB^ mice at 30 days, although not at 60 days, following 3-MCA injection (Fig. 4b, right panel). Early angiogenesis can temporarily enhance immune cell infiltration, and the irrigation within the tumour provide better oxygen and pH conditions for the cytotoxic activity of CD8^+^ T cells^69^. These results may explain our observed increase in immune cell infiltration in BL/6^NZB^ mice during early tumour progression (Fig. 1e-f). Further supporting our hypothesis, BL/6^NZB^ mice showed an increase in the measured CD31^+^ area in both chemically induced fibrosarcoma (Fig. 4c) and B16-F10 melanoma (Fig. 4d and Extended Data Fig. 4c) models. While these data indicate that tumour angiogenesis depends on the specific mtDNA variant, they also reveal an interplay with the immune system. Following BM transplantation (Fig. 2c), BL/6^NZB^ recipient mice with immune cells harbouring NZB mtDNA demonstrated an increase in CD31^+^ cells compared to their C57 counterparts (Extended Data Fig. 4b-c). Building on these findings and given that we previously showed that mROS induces TIL activation through PD-1 expression (Fig. 3), we next investigated whether the strain-specific differences in angiogenesis might also be linked to mROS generation. CD31^+^ cells from B16-F10 melanoma tumours harvested from BL/6^NZB^ and BL/6^C57^ mice showed no differences in total ROS levels between strains (Extended Data Fig. 4d). Furthermore, tumour vascularisation remained unchanged in our constitutive mCat^+^ mouse models (Fig. 4e and Extended Data Fig. 4e).

Finally, to address the cell specificity of this effect, we considered that H_2_O_2_ is highly permeable and could alter ROS levels in neighbouring cell types within the tumour microenvironment, potentially confounding tumour endothelial cell (TEC)-intrinsic signalling. For this reason, we generated a new mouse model expressing mCAT under the Cdh5-cre-ERT2 promoter (BL/6^NZB^ ECmCAT^+^) (Fig. 4f). This allowed us to directly assess whether TECs-derived mROS contributed to the higher angiogenic capacity in BL/6^NZB^ mice. We validated this model by observing reduced ROS levels in CD31^+^ cells in BL/6^NZB^ ECmCat^+^ mice compared to control BL6^NZB^ mice (Extended Data Fig. 4f-g). Our results showed no differences in B16-F10 melanoma tumour growth between BL6^NZB^ and BL6^NZB^ ECmCat^+^ mice (Fig. 4g). Furthermore, no significant differences were observed in tumour vascular density (Fig. 4h), in the total number of TILs (Extended Data Fig. 4h), in the frequency of tumour-infiltrating CD8^+^ T cells (Extended Data Fig. 4i), or in the effector CD8^+^PD-1^+^ population (Extended Data Fig. 4j).

In summary, BL/6^NZB^ mice establish a more favourable early tumoral vascular environment, potentially enabling greater lymphocyte infiltration and thereby improving tumour control. Moreover, the enhanced angiogenic capacity associated with the NZB mtDNA variant reflects a stromal property that is independent of mROS levels in tumour endothelial cells.

### BL/6^NZB^ mice exhibit enhanced CD8⁺ T cell-mediated antitumour responses and improved sensitivity to anti-PD-1 therapy

To gain deeper insight into the immune determinants and cell-specific contributors to the superior tumour control observed in BL/6^NZB^ mice, we first analysed the composition of the tumour microenvironment across strains. FACS analyses of B16-F10 melanoma tumours from BL/6^C57^ and BL/6^NZB^ mice revealed decreased frequencies of monocytes and neutrophils, along with increased numbers of conventional dendritic cells (cDCs) (Extended Data Fig. 5a-c). Although these alterations in monocytes, neutrophils and cDCs suggested a broader remodelling of the tumour microenvironment, CD8^+^ T cells remained the most likely mediators of the anti-tumour phenotype given our initial observations of enhanced T-cell infiltration and early activation signatures in BL/6^NZB^ tumours (Fig. 2). Next, we assessed the specific contribution of CD8⁺ T cells by performing adoptive CD8⁺ T-cell transfer (ACT) (Fig. 5a). *Rag1^⁻/⁻^* immunodeficient mice were used as hosts and received purified CD8^+^ T cells isolated from either BL/6^C57^ or BL/6^NZB^ donors, followed by subcutaneous injection of B16-F10 cells. Tumour growth was subsequently monitored, revealing that *Rag1^⁻/⁻^* mice receiving CD8^+^ T cells harbouring NZB mtDNA developed significantly smaller tumours (Fig. 5b). Although intratumoral CD8^+^ T frequencies did not differ significantly between groups (Fig. 5c), CD8^+^ T cells harbouring NZB mtDNA showed a higher proportion PD-1-expressing cells (Fig. 5d).

**Figure 5.**
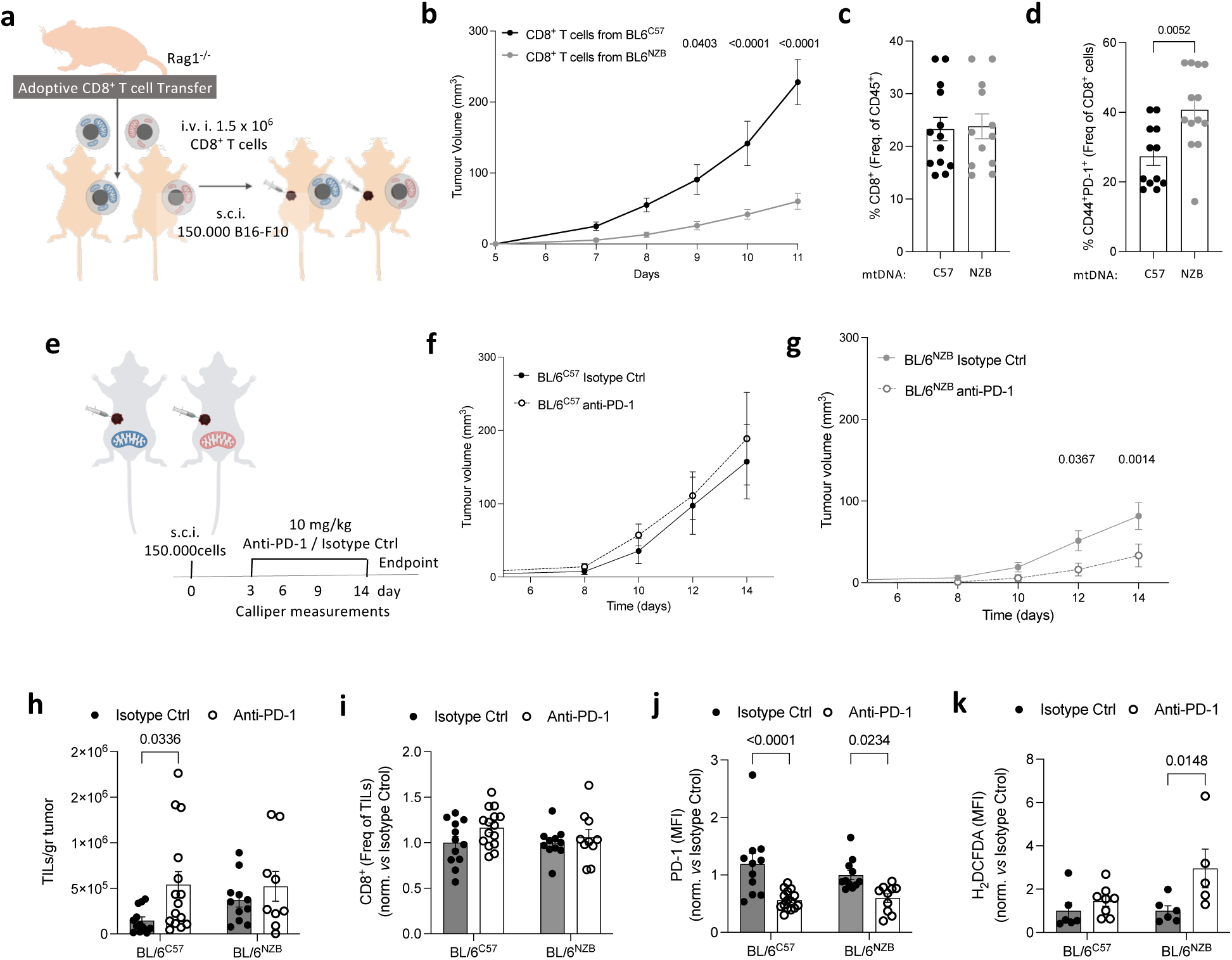
Tumour growth control is CD8⁺ T-cell dependent in BL/6^NZB^ conplastic mice, which show improved responsiveness to immune checkpoint inhibition. **a,** Schematic representation of the adoptive CD8^+^ T cell transfer experimental design. *Rag1^-/-^*mice lacking mature T and B cells, were intravenously injected (i.v.i.) with CD8^+^ T cells harbouring either C57 or NZB mtDNA. One day later, mice were subcutaneously injected (s.c.i.) with B16-F10 melanoma cells. **b,** Tumour growth measured at the indicated time points (n = 12 *Rag1^-/-^*mice injected with C57 mtDNA CD8^+^ T cells; n= 13 *Rag1^-/-^* mice injected with NZB mtDNA CD8^+^ T cells; data from two independent experiments). Two-way ANOVA with Fisher’s LSD test. **c-d,** FACS analysis of TILs in *Rag1⁻/⁻*mice bearing B16-F10 melanoma tumours, showing the percentage of CD8⁺ T cells among total CD45⁺ immune cells **(c)** and the percentage of PD-1⁺ effector CD8⁺ T cells among total CD8⁺ T cells **(d)** (n = 6 *Rag1⁻/⁻* mice injected with C57 mtDNA CD8⁺ T cells; n = 6 *Rag1⁻/⁻* mice injected with NZB mtDNA CD8⁺ T cells; representative of two independent experiments). Two-tailed unpaired t-test. **e,** Schematic representation of PD-1 treatment experimental design. Groups of BL/6^C57^ and BL/6^NZB^ mice were subcutaneously injected (s.c.i.) with B16-F10 melanoma cells, and anti-PD-1 antibody or isotype control was administered intraperitoneally (i.p., 50 mg/kg) on days 3, 6, and 9 post-injection. **f,** Tumour growth measured at the indicated time points in BL/6^C57^ mice (n=16 mice per treatment group; two independent experiments). **g,** Tumour growth measured at the indicated time points in BL/6^NZB^ mice (n=16 mice per treatment group; two independent experiments). Two-way ANOVA with Šídák’s multiple comparisons test. **h,** FACS analysis of TIL number normalised to tumour weight (n= 12 BL/6^C57^ Isotype Ctrl, n= 15 BL/6^C57^ anti-PD-1, n= 11 BL/6^NZB^ Isotype Ctrl, n= 9 BL/6^NZB^ anti-PD-1; two independent experiments). Two-way ANOVA with Šídák’s multiple comparisons test. **i,** FACS analysis showing the percentage of CD8^+^ T cells among total TCRβ^+^ cells (TILs). (n= 12 BL/6^C57^ Isotype Ctrl, n=15 BL/6^C57^ anti-PD-1, n= 11 BL/6^NZB^ Isotype Ctrl, n= 10 BL/6^NZB^ anti-PD-1; two independent experiments). Two-way ANOVA with Šídák’s multiple comparisons test. **j,** FACS analysis showing PD-1 expression (MFI) in CD8^+^ T cells from the indicated experimental groups. (n= 12 BL/6^C57^ Isotype Ctrl, n=15 BL/6^C57^ anti-PD-1, n= 11 BL/6^NZB^ Isotype Ctrl, n= 10 BL/6^NZB^ anti-PD-1; two independent experiments). Two-way ANOVA with Šídák’s multiple comparisons test. **k,** FACS analysis of cellular ROS levels in CD8^+^ T cells using H_2_DCFDA staining 14 days after subcutaneous injection of B16-F10 melanoma cells (n= 6 BL/6^C57^ Isotype Ctrl, n= 8 BL/6^C57^ anti-PD-1, n= 6 BL/6^NZB^ Isotype Ctrl, and n= 5 BL/6^NZB^ anti-PD-1). Two-way ANOVA with Šídák’s multiple comparisons test. Each dot represents an animal unless otherwise noted. Data are given as means ± SEM. *p-value < 0.05, **p-value < 0.01, and ****p-value < 0.0001.

These findings prompted us to speculate that mtDNA variation might influence the efficacy of clinically used immunotherapies based on immune checkpoint blockade. Among these, PD-1 inhibitors act by blocking the interaction between PD-1 and its ligand PD-L1 to restore T-cell activity^70^. To test this hypothesis, mice were treated with anti-PD-1 or isotype control antibodies administered intraperitoneally on days 3, 6, and 9 following B16-F10 tumour injection (Fig. 5e). Mice carrying NZB mtDNA showed a lower incidence of palpable tumours on day 10 after injection following anti-PD-1 treatment than all other experimental groups (Extended Data Fig. 5d). Tumour growth was monitored (Fig. 5f-g), and the analysis confirmed our previous results showing that tumours derived from BL/6^NZB^ mice had a lower volume increase compared to BL/6^C57^ mice (Fig. 1d and Fig. 5 f-g). Furthermore, our data indicate that the anti-PD-1 treatment strategy was more effective in BL/6^NZB^ mice (Fig. 5f-g). At the study endpoint, tumours were collected and TILs were subsequently analysed by FACS (Fig. 5h-k). The total amount of TILs was increased only in BL/6^C57^-derived tumours following anti-PD-1 treatment (Fig. 5h). Moreover, we confirmed that the treatment acted as expected: in both strains, anti-PD-1 did not alter the frequency of CD8^+^ T cells within the lymphocytic compartment of the tumour (Fig. 5i), while only the PD-1 surface marker on T cells was downregulated following treatment (Fig. 5j and Extended Data Fig. 5e). Lastly, given our earlier findings that CD8^+^ T cells with different mitochondrial genotypes display different ROS levels, and that increased mROS in these cells contributes to tumour control (Fig. 3), we reasoned that PD-1 blockade could further modulate their redox profile. We therefore assessed whether the decrease in PD-1 expression following treatment was accompanied by an inverse shift in intracellular ROS levels within CD8⁺ T cells (Fig. 5k and Extended Data Fig. 5f) and CD4⁺ T cells (Extended Data Fig. 5 g). Intracellular ROS levels in TILs increased more markedly in BL/6^NZB^ mice following anti-PD-1 treatment (Fig. 5k and Extended Data 5f-g).

Together, these findings demonstrate that cell-specific mtDNA variation in CD8⁺ T cells influences tumour control in mice. Moreover, anti-PD-1 therapy induced a more pronounced effect on both tumour onset and progression in mice harbouring NZB mtDNA, highlighting the functional relevance of mitochondrial genotype in shaping responsiveness to immune checkpoint blockade.

## Discussion

Mitochondria are active drivers and modulators of tumour development and progression. Variants and haplotypes in mtDNA add an additional layer of complexity, linking nucleo-mitochondrial interactions to cancer predisposition, progression, and therapeutic response. Here, we show that mitochondrial haplotypes impact on cancer generation and progression. Conplastic mice with NZB mtDNA developed fewer tumours and displayed slower tumour growth in both chemical and orthotopic models. This observation confirms and extend previous data^42^, and clearly points out a role for mtDNA in tumour generation and progression. This effect arose through distinct mechanisms, shaping the anti-tumour immune response both qualitatively and quantitatively in an mROS-dependent manner, and we showed that it was mediated by changes in TIL function. Moreover, we have further identified a stromal component contributing to mtDNA-dependent tumour control through angiogenesis. This appears largely independent of endothelial mROS, pointing to coordinated stromal-immune interactions rather than endothelial-autonomous mechanisms. Lastly, we demonstrate that mice carrying different mitochondrial haplotypes show differential responsiveness to anti-PD-1 treatment. Together, these data identify physiological mtDNA variants as key modulators of anti-tumour immunity and checkpoint inhibitor response.

Previous studies elegantly showed how tumour mitochondria could influence tumour invasiveness^22,23^ and the tumour microenvironment^71,72^. Importantly, the use of conplastic strains and transmitochondrial cybrids allows us to uncouple tumour-intrinsic from host-intrinsic mtDNA effects, providing genetic evidence that common mtDNA polymorphisms in the host compartment can shape tumour behaviour independently of the mtDNA carried by cancer cells themselves. Our finding that NZB mtDNA favours a PD-1^+^ effector CD8^+^ TIL phenotype fits with the concept that increased mitochondrial fitness and ROS-dependent signalling support robust T cell effector differentiation^45,73,74^. On the other hand, this could be due to a different ability of DCs to cross prime and induce a T cell response or to other innate cells. We also observed by BM transplant that non-immune components contribute to the phenotype, as highlighted by differences in neo-vasculature formation. This aligns with the concept that a transiently “normalised” vasculature can improve immune cell access and function within tumours and suggests that mtDNA haplotype may tune this window of favourable perfusion.

Immune-checkpoint inhibitors targeting CTLA-4, PD-1 or PD-L1 have transformed cancer therapy, yet benefit only a subset of patients^75,76^. Predictive biomarker analyses have linked treatment success to mitochondrial activation^77^, underscoring the relevance of our findings. Integrating our findings with recent enhancer-resolved maps of T-cell fate in tumours^78^ suggests that mtDNA-dependent ROS signalling may bias transcriptional trajectories towards effector or tissue-resident-like states. Mitochondrial features (including mtDNA mutations, respiratory complex composition and mitochondrial proteome remodelling^79–81^) can strongly influence tumour immunogenicity, immune evasion and clinical behaviour. Associations between mtDNA variation and cancer risk or immunotherapy response have been mainly limited by retrospective designs and geographically restricted cohorts^14,82–84^. Two recent studies highlight tumour-intrinsic mitochondrial dysfunction as a driver of immunotherapy outcome: *Mahmood et al.* demonstrated that truncating mtDNA mutations in complex I render melanoma cells more immunogenic and responsive to PD-1 blockade^27^; *Monson et al.* revealed that tumour-intrinsic defects in mitochondrial oxidative phosphorylation reprogramme metabolic signalling, generating a T-cell-excluded, immunosuppressive microenvironment that confers resistance to immune-checkpoint blockade^26^. Extending this framework beyond tumour-intrinsic alterations, our findings establish germline mtDNA as a fundamental and previously unrecognised determinant of mitochondrial function that operates via the immune system to preprogramme immune-cell metabolic sensitivity. Importantly, our analysis of the CheckMate-067 human cohort did not support a major role for European mtDNA macro-haplogroups in melanoma predisposition, indicating that the effects we observe are unlikely to reflect differential melanoma susceptibility across haplogroups. Instead, they point to mtDNA-dependent modulation of tumour immunity and therapeutic response as the primary mechanism underlying the observed differences.

Our results suggest that the mtDNA haplotype establishes a basal mitochondrial redox “set point” in T cells that determines whether PD-1-associated signalling is supported or disrupted by changes in mROS. In BL/6^C57^ mice, which exhibit lower basal mROS, further depletion of mitochondrial H_2_O_2_ by mCat may push T cells below a signalling threshold required to sustain PD-1 expression and effector programming, thereby weakening anti-tumour function and accelerating melanoma progression. By contrast, in BL/6^NZB^ mice with higher basal mROS, mCat-mediated scavenging may reduce excess mROS without dropping below this functional window, resulting in minimal changes in PD-1 levels and TIL phenotype. Host mtDNA haplotype alone can tune CD8^+^ T-cell redox state, PD-1 expression and effector function, thereby shaping the magnitude of response to checkpoint inhibition. PD-1 engagement increases cellular ROS and constrains oxidative metabolism in T cells, and that PD-1 blockade can reduce ROS and alter susceptibility to metabolic stress^6^. This apparent discrepancy highlights that PD-1/ROS coupling is context dependent and may differ between alloreactive versus tumour-specific T cells, between total ROS and compartmentalised mROS, and across mitochondrial genotypes. Together, these findings highlight an emerging therapeutic landscape in which tuning mitochondrial ROS, rather than global cellular ROS, offers a mechanistically precise strategy to boost antitumour immunity and improve responses to immunotherapy. The magnitude of these effects are likely haplogroup dependent, reflecting differences in basal mitochondrial ROS production^40^, and may likewise vary across tumour contexts^74,85–87^.

The emerging landscape of mitochondrial immune-oncology shows that mitochondrial crosstalk between tumour cells and T cells, through transfer of mutant mtDNA or selective defects in the electron transport chain, can powerfully shape antitumour immunity and dictate responsiveness to checkpoint blockade^79,80^. Mitochondrial features are now being explored as therapeutic avenues, from boosting T cell bioenergetics to targeting neoantigens derived from mitochondrial mutations^88,89^. Our findings extend this rapidly evolving field by demonstrating that common germline mtDNA variation in the host can reprogramme tumour angiogenesis, TIL infiltration, PD-1 expression and mROS signalling. These findings suggest opportunities to fine tune T cell mitochondrial function and the mitochondrial ROS signals that shape immunity, thereby improving responses to immunotherapy.

## Material and Methods

### Lead Contact and Materials Availability

Further information and requests for resources and reagents should be directed to and will be fulfilled by the Lead Contacts, Jose Antonio Enríquez and Ana Victoria Lechuga-Vieco.

### Data and code availability

All data reported in this paper will be shared by the lead contact upon request. This paper does report original code. Any additional information required to reanalyse the data presented in this work is available from the Lead Contact upon request.

### Experimental model and subject details

#### Animals

All animal procedures conformed to EU Directive 86/609/EEC and Recommendation 2007/526/EC regarding the protection of animals used for experimental and other scientific purposes, enforced in Spanish law under Real Decreto 1201/2005. Animal studies have been conducted in accordance with, and with the approval of, the United Kingdom Home Office. All procedures were done under the authority of the appropriate personal and project licenses issued by the United Kingdom Home Office License number PBA43A2E4 or approved by the Animal Subjects Committee of the Instituto de Salud Carlos III (Madrid, Spain) in accordance with EU Directive 86/609/EEC and Madrid Community Organs in the PROEX 98.6/20, PROEX 026/17 and PROEX 172/19. Approval of the different experimental protocols required estimation of minimum sample size and definition of the randomization and blinding criteria. Mice were housed in the local animal facility under specific pathogen–free conditions, a 12-hour light/dark cycle and had ad libitum access to water and a standard chow diet (5K67 LabDiet). The conplastic mouse strains with C57BL/6 nuclear genome and either C57 or NZB/OlaHsd mtDNA were described before^42^. B6-SJL T mice expressing the CD45.1 allele (Ptprca Pepcb/BoyJ), OT-I transgenic mice (C57BL/6-Tg (TcraTcrb)1100Mjb/J), tamoxifen-inducible cre/ERT2 mice in endothelial cells (Tg(Cdh5-cre/ERT2)1Rha) and *Rag1^-/-^*mice (B6.129S7-Rag1tm1Mom/J) were purchased from The Jackson Laboratory (Bar Harbor, ME, USA). Generation of mCat^+^ mice was described in^66^.

#### Human samples

We analysed UK Biobank (UKB) data^48^, which reflect well-established patterns in the UK population. Additionally, observed frequencies were derived from trial participants in NCT01844505 (Creative Commons Attribution 4.0 International License)^26^.

### Methods Details

#### Macro-haplogroup bias analyses

To assess potential enrichment or bias in macro-haplogroup representation within the CheckMate-067 clinical trial population, we performed a binomial test for each macro-haplogroup of interest (H, U, K, T, J, and a composite group comprising W, V, R, I, and X). Expected distributions were defined using macro-haplogroup frequencies derived from UK Biobank (UKB) data^48^, which reflect well-established patterns in the UK population. The observed frequencies among trial participants (NCT01844505)^26^ were then compared to these expectations to determine if significant deviations occurred.

#### *In vitro* B16-F10 cell culture

B16-F10 melanoma cells were cultured in Dulbecco’s Modified Eagle Medium (DMEM; Sigma-Aldrich, D6429) supplemented with 10% foetal bovine serum (FBS; Gibco, A5256801), 1 mM penicillin–streptomycin (Gibco, 15140-122), 1 mM L-glutamine (Gibco, 25030-024) and 1 mM sodium pyruvate (Gibco, 11360070). Cultures were maintained at 37 °C in a humidified incubator with 5% CO₂. For passaging and preparation for *in vivo* injections, cells were detached using 0.25% trypsin-EDTA (ThermoFisher Scientific, 25200072). Cell pellets were then resuspended in sterile PBS and adjusted to the required concentration for subcutaneous injection in mice for tumour induction.

#### Generation of transmitochondrial cell lines

(a) For mitochondrial genome repopulation, cultures were treated with 3 μg/mL rhodamine 6G (R6G; Sigma, R4127) in DMEM supplemented with 5% FBS, 50 μg/mL uridine, and 1% antibiotic for 72-96 hours, refreshing the medium each 24 hours. Two hours prior to fusion, the culture medium was exchanged for DMEM containing 5% FBS, 50 μg/ml uridine (Sigma-Aldrich, U3750), and 1% Penicillin-Streptomycin.

(b) Enucleation of donor cells was performed by treating confluent monolayers with 1 μg/mL cytochalasin B and mitomycin for 1 hour, followed by centrifugation at 6,500-7,200 x r.p.m. for 20 min at 37 °C to remove nuclei.

Enucleated cells (b) were fused with recipient cells (a) in the presence of 40-50% polyethylene glycol (PEG; Serva 33123), mixing for 60 seconds. Cells were washed with DMEM containing 10% DMSO and plated in DMEM containing 5% FBS, 50 μg/ml uridine, and 1% antibiotic. Medium was replaced every 2 days for the first week, after which they were expanded under standard uridine-free medium.

#### Spontaneous tumour development assessment

Survival studies were performed, and mice found dead during daily inspections were recorded as censored deaths and necropsied for tumour inspection. Animals meeting endpoint criteria were excluded from the analysis in long-term experiments. Tumours identified at necropsy were collected for histopathological examination to determine tumour type and confirm gross morphological observations.

#### 3-MCA fibrosarcoma generation

8-12-weeks-old BL/6^C57^ and BL/6^NZB^ mice received a single intra-muscular injection of 40 μl solution containing 3-methyl-cholanthrene (3-MCA) (Sigma-Aldrich) in one of the rear legs. 3-MCA was used at a concentration of 25 μg/μl and dissolved in sesame oil (Sigma). Mice were monitored regularly and sacrificed 90 or 120 days after the injection.

#### Primary melanoma tumour generation

Mice were injected subcutaneously into the right flank with 1.5 x 10^5^ B16-F10 or B16-F10-OVA cells in 0.1 ml PBS for immunogenicity or tumour growth experiments. Tumour size was measured every two days from day 8 to day 14 post-injection using a digital calliper. Tumour volumes were calculated as follows: V = 1/2 × (L × M^2^), where L represents the long axis and M the short axis of the tumour. On day 14, tumours were excised for downstream analyses.

*Adoptive T cell transfer experiments*. *Rag1^−/−^*mice (B6.129S7-Rag1tm1Mom/J) were used as recipients for adoptive T cell transfer. Total CD8^+^ T cells were isolated from splenocytes by immunomagnetic negative selection according to manufacturer’s instructions (Stemcell, 19853) using EasyEights^TM^ EasySep^TM^ Magnet (StemCell, 18103). Purified CD8^+^ T cells carrying either C57 or NZB mtDNA were intravenously injected into *Rag1^−/−^* recipients (1.5 × 10^6^ cells per mouse). One day later, mice were subcutaneously injected in the right flank with 1.5 × 10^5^ B16-F10 melanoma cells.

*PD-1 Blockade Treatment.* Anti–PD-1 antibody (clone RMP1-14, BioXCell, Cat. No. BE0146) was administered intraperitoneally at 10 mg/kg on days 3, 6, and 9 after tumour induction. Control animals received the same dosing schedule of rat IgG2a isotype control (clone 2A3, BioXCell, Cat. No. BE0089).

*N-acetyl-L-cysteine (NAC) treatment.* Mice were provided with drinking water containing 61.2 mM N-acetyl-L-cysteine (NAC; Sigma, Cat. No. A9165), corresponding to approximately 1 % (w/v), beginning 10 days prior to experimental procedures. NAC-containing water was supplemented with orange flavouring to facilitate palatability and was protected from light throughout the treatment period. Solutions were replaced twice weekly and maintained until the end of the experiment.

*MitoQ treatment.* Mice received intraperitoneal (i.p.) injections of MitoQ (15 mg/kg; provided by Dr. M. Murphy) or vehicle control (PBS; in the flak opposite to the tumour). The first dose was administered 2 days prior to melanoma induction, followed by repeated injections every 2 days throughout the tumour growth period.

#### Bone marrow chimaeras

8–12-week-old BL/6^C57^ and BL/6^NZB^ mice were lethally irradiated with a total dose of 11 Gy (550 cGy twice, 4 h apart) and rested for 1 h prior to transplantation. A total of 1.5 × 10^6^ bone marrow (BM) cells from 8–12-week-old BL/6^C57^ or BL/6^NZB^ donor mice were injected intravenously into the irradiated recipients. Reconstitution efficiency was monitored at 2-, 4-, 6-, and 8-weeks post-irradiation by haematological analysis of peripheral blood. For this, 50 μl of blood was collected from the tail vein into Microvette® 100 EDTA tubes (Sarstedt) and kept at 4 °C until analysis. Complete blood counts were performed using a Pentra 80 automated haematology analyser. Tumour induction experiments were performed 8–10 weeks after irradiation, once stable haematopoietic reconstitution was confirmed.

#### *In vivo* magnetic resonance imaging (MRI)

*In vivo* MRI in mice was performed with an 7T horizontal Agilent/Varian scanner (Agilent, Santa Clara, USA) equipped with a DD2 console and an active-shielded 205/120 gradient insert coil with 130 mT/m maximum gradient strength and a combination of volume coil/two channel phased-array (Rapid Biomedical GmbH, Rimpar, Germany). Mice were anaesthetised with 2% isoflurane (Abbott) and oxygen and positioned on a thermoregulated (35-37°C) mouse bed with continuous monitoring of the respiratory cycle. Ophthalmic gel was placed in the eyes to prevent retinal drying. Three-dimensional gradient-echo volumetric imaging was performed with minimum repetition and echo times (TR/TE = 3 ms/1.5 ms), 20° flip angle, isotropic 150 µm spatial resolution, and a 6 cm × 6 cm field of view (FOV), totalling ∼6 min acquisition time. For 2D axial acquisitions, consecutive 1 mm-thick slices were acquired using a gradient echo sequence with TR/TE = 40 ms/4 ms, 20° flip angle, and 100 kHz bandwidth, for a total scan time of 80 s. Images were acquired after intravenous administration of gadopentetate dimeglumine (Magnevist, 2 mM; Bayer Healthcare Pharmaceuticals) via tail vein injection. Tumour volumes were quantified using OsiriX software (Pixmeo) from the acquired 3D datasets. Following euthanasia, tumours were excised, weighed, and measured with a calliper. Representative axial slices from the 3D volume reconstructions are shown in Extended Data Fig. 1.

#### Directed *in vivo* angiogenesis assay (DIVAA)

Angiogenic responses were assessed using the manufacturer’s protocol (Cultrex, 3450-048-IK) and quantified as previously described^90^.

#### Immunohistochemistry (IHC)

Tumours were collected from BL/6^C57^ and BL/6^NZB^ mice, fixed with 4% PFA for 24 hours and sectioned at 5 μm. Three sections from each tumour at three different levels of the tissue (80-100 μm between each section was discarded) were processed, mounted on a glass slide, stained with Ki67 (Invitrogen PA5-19462), CD3 (Invitrogen, MA1-90582) and CD31 (Abcam, ab28364) primary antibodies and HRP polymer conjugated secondary antibodies. Slides were digitalized using NDP view2 Software and quantification of the staining was performed using ImageJ^91^.

#### *Ex vivo* B16-F10 tumour growth

Tumour samples were processed in 24-well plates, then incubated for 15 min at 37 °C in 1 mL of digestion buffer containing 2 mg/mL collagenase D (Sigma-Aldrich, 1108886601) and 1 mg/mL DNase I (Sigma-Aldrich, EN0521) in RPMI-1640 (Sigma-Aldrich, R8758-500). After the first incubation, the tissue suspension was pipetted up and down repeatedly to further dissociate the material, followed by a second 15 min incubation at 37° C. The resulting single-cell suspension was transferred to p150 tissue culture plates (Fisher Scientific, 11804125), where cells were maintained under standard conditions. Adherent melanoma cells were permitted to attach and proliferate, whereas non-cancerous cells were gradually eliminated through successive medium changes and passaging. Total cell numbers were determined using a Neubauer haemocytometer by mixing cell suspension with trypan blue

#### Flow cytometry phenotyping

Tumour samples were pierced and minced in 24-well plates and incubated for 15 minutes at 37°C in 1 mL of digestion buffer (2 mg/mL collagenase D, and 1 mg/mL DNAse I in RPMI-1640, both from Sigma-Aldrich). After the first 15 minutes of incubation, cells were pipetted up and down repeatedly, then returned for a second 15-minute incubation at 37° C. After digestion, B16 cells were stained with H-2Kb (clone AF6-88.5.5.3), CD105 (clone MJ7/18) and PD-L1 (clone 10F.9G2) specific antibodies. T cells from the tumour were stained with CD8 (clone 53-6.7), PD-1 (clone 29F.1A12), CD44 (clone IM7), TCR β (clone H57-597), CD62L (clone MEL-14) (all from biolegend), CD25 (clone 3C7) and FOXP3 (clone MF-14). Myeloid cells from the tumour were stained with CD45–APCefluor780 (30-F11), F4/80–PerCP-Cy5.5 (BM8), CD103–BV421 (M290 (RUO)), Ly6G–APC (1A8 (RUO)), XCR1–PE (ZET), CD24–PE-Cy7 (M1/69) Aqua Live/Dead Pacific Orange, B220–BV786 (RA3-6B2 (RUO)), Ly6C–PerCP-Cy5.5 (HK1.4), CD11b–PE-Cy7 (M1/70), CD11c–BV650 (N418), MHCII–A700 (M5/114.15.2). Cells were stained with Live/Dead fixable dye according to manufacturer instruction (Thermo Fisher). For intracellular marker staining samples were fixed permeabilized with foxp3 staining kit (Life Technologies, 00-5523-00) according to manufacturer instruction. Samples were acquired on a FACS Fortessa (BD Biosciences) flow cytometer, and data were analysed with FlowJo version 10.4.1. Compensation beads (eBioscience) were used to generate the compensation matrix, and fluorescence minus one (FMO) were used as control.

*ROS measurements.* Reactive oxygen species (ROS) levels were measured in tumour endothelial cells (TECs) and tumour infiltrating lymphocytes (TILs) using 2’,7’-dichlorodihydrofluorescein diacetate (H_2_DCFDA, ThermoFisher, 0.4mM final concentration), and MitoSOX™ Red (ThermoFisher, 5 μM final concentration). A minimum of 2000 events were recorded for each sample using the BD LSR II flow cytometer (BD Biosciences). All experiments were performed in triplicate. Samples were analysed and the mean fluorescence intensity (MFI) was calculated using FlowJo Software (v10.7.1).

#### Tissue processing and immunofluorescence staining

Mice were euthanized by CO_2_ inhalation, and tumours were carefully dissected and fixed overnight at 4° C in 4% paraformaldehyde (PFA) prepared in PBS. Following fixation, tissues were washed three times in PBS and cryoprotected overnight at 4° C in 30% sucrose (Sigma) in PBS. Samples were embedded in OCT^TM^ compound (Sakura) and stored at −80° C until sectioning. Cryosections (20 µm) were obtained using a Leica cryostat and washed three times in PBS (10 min each). Sections were then blocked and permeabilized in PBS containing 10% donkey serum (Millipore) and 1% Triton X-100 for 1 h at room temperature. Primary antibodies were diluted in the blocking/permeabilization buffer and incubated overnight at 4° C. After three 10 min washes in PBS, sections were incubated for 2 h at room temperature with species-appropriate fluorophore-conjugated secondary antibodies (1:400, Jackson ImmunoResearch) and DAPI for nuclear counterstaining. Following three final PBS washes, sections were mounted in Fluoromount-G (SouthernBiotech). The following primary antibodies were used: purified rat anti-mouse CD31 (1:400, Pharmingen, 553370), rabbit anti-ERG-Alexa647 (1:200, Abcam, ab110639), monoclonal rat anti-Endomucin (1:400, Santa Cruz Biotechnology, sc-53941), and rat anti-mouse VE-cadherin clone 11D4 (1:200, BD Biosciences, 555289).

*Confocal imaging and vascular quantification.* Tumour sections were imaged at high resolution using a Leica SP8 confocal microscope with ×20 or ×40 objectives. Individual fields or tiled images of large areas were acquired. Image analysis was performed in Fiji/ImageJ^91^, where automatic thresholding was applied to select and quantify object areas. Autofluorescence signal was used to define the total tumour area, while CD31, Endomucin, or VE-Cadherin signals were used to determine vascularized areas, and ERG staining was used to quantify TEC numbers. The following vascular parameters were calculated: (i) Vascular density = 100 × (vascularized area / tumour area); (ii) Endothelial density = 100 × (endothelial cell number / vascularized area)

## Statistical analysis

All data are presented as the mean ± standard error of the mean (SEM). For comparisons between two groups, unpaired two-tailed Student’s t-tests were used for normally distributed datasets with equal variances; otherwise, a two-tailed t-test was applied as appropriate. For datasets with ≥3 groups, one-way analysis of variance (ANOVA) followed by Tukey’s multiple-comparison test was performed. Paired or unpaired one-way ANOVA was used for multiple comparisons of normally distributed datasets with one variable. Statistical significance was determined based on the p-value for the hypothesis being tested. Analyses were carried out in GraphPad Prism (version 9.5.1; GraphPad Software, San Diego, CA, USA).

## Acknowledgements

We acknowledge all the members of the Cerundolo and Lechuga-Vieco research groups for their insightful scientific discussions and contributions to this work. We thank Prof. Juan P. Bolaños for kindly providing the mitochondria-targeted catalase mouse model, and Prof. Mike Murphy for providing MitoQ. We thank Richard Corderoy from the QQF, BMS at University of Oxford, CNIC and the Barcelona Science Park (PCB) animal facility for their excellent care and assistance of animal well-being. WIMM and CNIC flow Cytometry and Microscopy Facilities. We thank Dr Sergio Caja and Eva Raquel Martínez Jiménez for their experimental assistance. Antonio de Molina-Iracheta from the CNIC Comparative Medicine Unit for his valuable input and discussions, and the CNIC Histopathology laboratory. Images were created using BioRender.com under the academic license granted to JAE.

This work was supported by: The”ReDIB ICTS infrastructure TRIMA@CNIC, Ministerio de Ciencia e Innovación (MCIN). Institutional core funding to the MRC Translational Immunology Discovery Unit, Radcliffe Department of Medicine, University of Oxford to GP and UG. Ministerio de Ciencia, Innovación y Universidades (MCIU) PhD Fellowship (PRE2019-087462) to RJ-M. MCIU PhD Fellowship (PRE2019-087459) and the Instituto de Salud Carlos III, AC23_2/00019 to MdA-L. This project has received funding from ISCIII and was co-funded by the European Union under the Horizon Europe Framework Programme No. 10105426. JLC-A is supported by Fundación “la Caixa”(LCF/PR/HR23/52430010). The European Union’s Horizon Europe programme through a Marie Skłodowska-Curie Action doctoral fellowships (Grant Agreement No. 101126676) to VL. Formación de Personal Universitario (FPU) PhD Fellowship (FPU16/03142) from the Spanish Ministry of Education to SCK. Ramón y Cajal Program (RYC2023-043157-I) funded by the MCIN/AEI/10.13039/501100011033 and the ESF“Investing in your future” to JP. The research in RB laboratory was supported by the European Research Council Consolidator Grant AngioUnrestUHD (101001814) and ‘la Caixa’ Banking Foundation MapVM (HR19-00120). JR-C is funded by grant PID2021-123238OB-I00 and PID2024-155807OB-C21 funded by MCIN/AEI/10.13039/501100011033 by”ERDF A way of making Europe and by the“European Union”, grant DTS24/00043 & ISCIII-AES-2024/000225 funded by Instituto de Salud Carlos III (ISCIII), Basque Government under the Elkartek 2024 Program (KK-2024-00041) and Elkartek 2025 (KK-2025-00084) and from the Fundación contra la Hipertensión Pulmonar (Empathy). Work in the DS laboratory is funded by the CNIC; by Ministerio de Ciencia, Innovación y Universidades (MICIU) PID2022-137712OB-I00, CPP2022-009762 and CPP2024-011365 MICIU/AEI/10.13039/501100011033 Agencia Estatal de Investigación, Unión Europea NextGenerationEU/PRTR; by Comunidad de Madrid (P2022/BMD-7333 INMUNOVAR-CM); by Scientific Foundation of the Spanish Association Against Cancer (AECC--PRYGN246642SANC); by Worldwide Cancer Research WWCR-25-0080; by European Union ERC-2023-PoC; by a research agreement with Inmunotek S.L.; and by “la Caixa” Foundation (LCF/PR/HR23/52430012 and LCF/PR/HR22/52420019). JAE is supported by PID2021-1279880B-100 and TED2024-158440OB-I00 funded by MCIN/AEI/10.13039/501100011033, the European Union “NextGenerationEU”/Plan de Recuperación Transformación y Resiliencia/PRTR, CIBERFES (CB16/10/00282) and ERC-2024-ADG (GA 101198761). The CNIC is supported by the Instituto de Salud Carlos III (ISCIII), the Ministerio de Ciencia, Innovación y Universidades (MICIU) and the Pro CNIC Foundation, and is a Severo Ochoa Center of Excellence (grant CEX2020-001041-S funded by MICIU/AEI/10.13039/501100011033). Fundación Alfonso Martín Escudero Foundation LT-postdoctoral Fellowship (FAME2019), “La Caixa” Foundation (ID 100010434, LCF/BQ/PI24/12040005), Domingo Martínez Foundation and the Spanish Association for Cancer Research (ASEICA) and funding from MINECO through the Centers of Excellence Severo Ochoa Award and CERCA Programme of the Generalitat de Catalunya to AVL-V.

We dedicate this study to the memory of the late Professor V. C., FRS, an inspirational colleague, mentor, and pioneer in the field of cancer immunology and immunotherapy, who passed away in January 2020.

## Author contributions

GP, UG, JAE and AVL-V conceptualized and designed the study. AVL-V, GP, and RJ-M performed and analysed experiments, methodology, and investigation. GP, UG, RJ-M, JAE and AVL-V discussed analyses. GP and AVL-V supervised the project and interpreted the experimental data. GP, RJ-M and AVL-V wrote the original draft. GP, RJ-M, UG, AVL-V and JAE reviewed and edited the manuscript. MdA-L and RB performed TEC analyses by confocal imaging to assess angiogenesis in *ex vivo* tumour samples. JLC-A analysed the prevalence of mitochondrial haplogroups in patient cohorts using UK Biobank Data. JP and JR-C conducted the *in vivo* imaging analyses. RJ-M, MMM-H, VL and RMdM performed and monitored tumour growth in mice. MMM-H carried out the DIVAA assay analyses. SCK and DS validated the findings using immunodeficient mouse models. All authors reviewed and approved the final manuscript.

---

**Extended Data Figure 1. Characterisation of tumour models in conplastic strains.**

**a,** Enrichment analysis for European Macro-Haplogroups in patients with advanced melanoma. In all cases, null hypothesis of random distribution of Macro-Haplogroups was evaluated applying a binomial test. In this analysis a threshold for adjusted p-value<0.05 corrected by Benjamini-Hochberg method.

**b,** Schematic diagram of 3-methylcholanthrene (3-MCA)-induced fibrosarcoma.

**c,** Representative coronal T1-weighted MRI sections acquired after intravenous injection of a gadolinium-based contrast agent. Tumours were imaged on day 60 after intramuscular injection (i.m.i.) of 3-MCA. Chemically induced tumours are labelled in red, and data were collected from 8-10 mice per mtDNA haplotype group.

**d,** Schematic representation of the orthotopic B16-F10 melanoma model experimental design.

**e,** Representative immunohistochemical images of tumour sections, collected on day 14 after subcutaneous injection (s.c.i.) of BL/6^C57^ and BL/6^NZB^ mice, stained for Ki-67 (left panel) and quantified based on the percentage of Ki-67^+^ tumoral area (right panel). Each dot represents a different tumour section (n= 13 BL/6^C57^ mice and n= 11 BL/6^NZB^ mice; two independent experiments). Scale bar: 250μm. Kolmogorov-Smirnov nonparametric test.

**f,** Tumour growth of the two strains following subcutaneous injection of a newly generated transmitochondrial cybrid B16-F10 cell line carrying NZB mtDNA (n=5-6 mice per host strain). Two-way ANOVA with Šídák’s multiple comparisons test.

**g,** Quantification of the CD45^+^ area in immunohistochemical sections of melanoma tumours from BL/6^C57^ and BL/6^NZB^ mice. Each dot represent*s* a different tumour section (n= 15 BL/6^NZB^ mice and n= 11 BL/6^NZB^ mice; two independent experiments). Kolmogorov-Smirnov nonparametric test.

Data are given as means ±SEM. *p-value < 0.05, and **p-value < 0.01.

**Extended Data Figure 2. Flow cytometry analysis of immune and tumour cell populations in B16-F10 melanoma tumours.**

**a-b,** Gating strategy used in FACS to identify CD8^+^CD44^+^PD-1^+^ subsets within TILs in 3-MCA fibrosarcomas (**a**) and B16-F10 melanoma tumours (**b**).

**c,** FACS analysis showing the percentage of CD4^+^ cells among total CD45^+^ immune cells in 3-MCA-induced fibrosarcomas (n= 5 mice per genotype).

**d,** FACS analysis showing the percentage of CD4^+^ cells among total CD45^+^ immune cells in B16-F10 melanoma tumours (left panel), and the percentage of effector CD4^+^ T cells expressing PD-1 within total CD3^+^ lymphocyte population (right panel) (n= 7 BL/6^C57^ and n= 6 BL/6^NZB^ mice). Data are representative of 3 independent experiments.

**e,** FACS analysis of PD-L1 expression in B16-F10 tumour cells excised from conplastic mice. PD-L1 expression was quantified in CD105^+^PD-L1^+^ cells (MFI, median fluorescence intensity; left panel) and representative FACS histograms (right panel) obtained from tumours derived from the indicated strains (n= 6 BL/6^C57^ and n= 8 BL/6^NZB^ mice). Two-tailed unpaired t test. Data are representative of 3 independent experiments.

**f,** Gating strategy used in FACS analysis to identify PD-L1-expressing cells within the CD105⁺ tumour cell population from B16-F10 tumours of the indicated genotypes

Each dot represents an animal unless otherwise noted. Data are given as means ±SEM. *p-value<0.05.

**Extended Data Figure 3. TILs and tumour cell characterisation in models expressing mitochondria-targeted catalase.**

**a,** FACS analysis of TILs showing the percentage of CD4^+^ cells in TILs (n=3-5 mice per group). One-way ANOVA test.

**b,** FACS analysis of total cellular ROS levels (H_2_DCFDA staining) and mROS levels (MitoSOX staining) in CD4^+^ T in B16-F10 melanoma tumours excised after 14 days of injection in control and mCat^+^ conplastic mice (n=6 mice per genotype). Two-way ANOVA with Šídák’s multiple comparisons test.

**c,** Representative immunohistochemical images of melanoma tumour sections from BL/6^C57^, BL/6^NZB^, BL/6^C57^ mCat^+^ and BL/6^NZB^ mCat^+^ mice, stained for CD3 (left panel), and quantified based on the percentage of CD3^+^ area (right panel), with each dot representing a different tumoral section (n=4-6 mice per genotype). Scale bar: 250μm.

**d,** FACS analysis showing the percentage of effector CD8^+^ T cells expressing PD-1 in TILs (n=6-8 per genotype). Two-way ANOVA with Šídák’s multiple comparisons test.

**e,** FACS analysis of PD-L1 expression (MFI) in B16-F10 tumour cells 14 days after subcutaneous injection (n = 6–8 mice per genotype). Two-way ANOVA with Šídák’s multiple comparisons test.

Each dot represents an animal unless otherwise noted. Data are given as means ±SEM. *p-value < 0.05.

**Extended Data Figure 4. Characterisation of endothelial, redox and immune features in B16-F10 melanoma tumours.**

**a,** Tumour weight of BL/6^C57^ and BL/6^NZB^ 14 days post injection of B16-F10 melanoma cells. 8–10 weeks prior to tumour challenge, mice were irradiated, and each strain was repopulated with the bone marrow from BL/6^C57^ or BL/6^NZB^ mice covering all the possible combinations (n=3-6 per group; two independent experiments). Two-way ANOVA with Šídák’s multiple comparisons test.

**b,** FACS gating strategy to identify CD31^+^ tumour cells within the CD105^+^ population in BL/6^NZB^ mice reconstituted with BL/6^C57^ (left panel) or BL/6^NZB^ (right panel) bone marrow.

**c,** FACS analysis showing percentage of CD31^+^ cells within total live cells 14 days post injection of B16-F10 melanoma cells in BM transplanted BL/6^C57^ and BL/6^NZB^ mice (n=4-6 per group). Two-way ANOVA with Šídák’s multiple comparisons test.

**d,** FACS analysis of cellular ROS levels in CD31^+^ cells using H_2_DCFDA staining 14 days post injection of B16-F10 melanoma cells in BL/6^C57^ and BL/6^NZB^ mice (n=6 mice per genotype). Two-tailed unpaired t test.

**e,** Representative immunohistochemical images of melanoma tumour sections from control and mCat^+^ conplastic mice, stained for CD31 (left panel) and, percentage of CD31^+^ area in tumours, with each dot representing a different tumoral section (right panel) (n= 9 BL/6^C57^, n= 8 BL/6^NZB^, n= 14 BL/6^C57^ mCat^+^ and n= 11 BL/6^NZB^ mCat^+^). Two-way ANOVA with Šídák’s multiple comparisons test. Scale bar: 250μm.

**f-g,** FACS analysis of cellular ROS levels in CD31^+^ cells using H_2_DCFDA staining 14 days post injection of B16-F10 melanoma cells showing H_2_DCFDA MFI (median fluorescence intensity) analysis (**f**) and, representative histograms of H_2_DCFDA in ECs obtained from tumours of the indicated genotypes (**g**) (n= 10 BL/6^NZB^ and n= 4 BL/6^NZB^ ECmCat^+^ mice). Two-tailed unpaired t test.

**h-j,** FACS analyses showing total TILs normalised to tumour weight (**h**), the proportion of CD8^+^ cells within CD45^+^ cells (**i**), and the percentage of PD-1^+^ effector CD8^+^ T cells (**j**), from BL/6^NZB^ mice and BL/6^NZB^ ECmCat^+^ mice 14 days after B16-F10 injection (n= 4–10 per genotype). Two-tailed unpaired t test.

Each dot represents an animal unless otherwise noted. Data are given as means ± SEM. *p-value < 0.05, **p-value < 0.01, and ****p-value < 0.0001.

**Extended Data Figure 5. FACS analysis of intratumoral immune populations. a,** Gating strategy used in FACS analysis to identify the principal myeloid populations, including conventional dendritic cell subsets (cDC1 and cDC2), tumour-associated macrophages (TAMs), monocytes, and neutrophils.

**b,** Representative flow cytometry plots for cDC1 and cDC2 in tumours from BL/6^C57^ (left) and BL/6^NZB^ (right) mice.

**c,** FACS analyses showing different intratumoral CD45^+^ cell subsets. From left to right: monocytes, neutrophils, TAMs, monocyte-derived dendritic cells (MoDCs), and cDCs (n=7 mice per genotype). Two-tailed unpaired t-test.

**d,** Frequency of mice with palpable tumours on day 10 after B16-F10 cell injection (n=16 mice per indicated experimental group; two independent experiments). Two-tailed unpaired t-test.

**e,** FACS analysis showing PD-1 expression (MFI) in CD4^+^ T cells from the indicated experimental groups (n= 6, BL/6^C57^ Isotype Ctrl, n= 8 BL/6^C57^ anti-PD-1, n= 6 BL/6^NZB^ Isotype Ctrl, and n= 6 BL/6^NZB^ anti-PD-1). Two-way ANOVA with Šídák’s multiple comparisons test.

**f,** Representative FACS histograms of intracellular ROS levels (H_2_DCFDA) in CD8^+^ T cells isolated from tumours of the indicated strains and conditions.

**g,** FACS analysis of cellular ROS levels in CD4^+^ T cells using H_2_DCFDA staining 14 days after subcutaneous injection of B16-F10 melanoma cells (n= 6 BL/6^C57^ Isotype Ctrl, n= 8 BL/6^C57^ anti-PD-1, n= 6 BL/6^NZB^ Isotype Ctrl, n= 6 BL/6^NZB^ anti-PD-1). Two-way ANOVA with Šídák’s multiple comparisons test.

Each dot represents an animal unless otherwise noted. Data are given as means ± SEM. *p-value < 0.05, **p-value < 0.01, and ****p-value < 0.0001.

