## Extended Data for "Mitochondrial genetics defines anti-tumour immunity through mitochondrial ROS and PD-1 signalling"

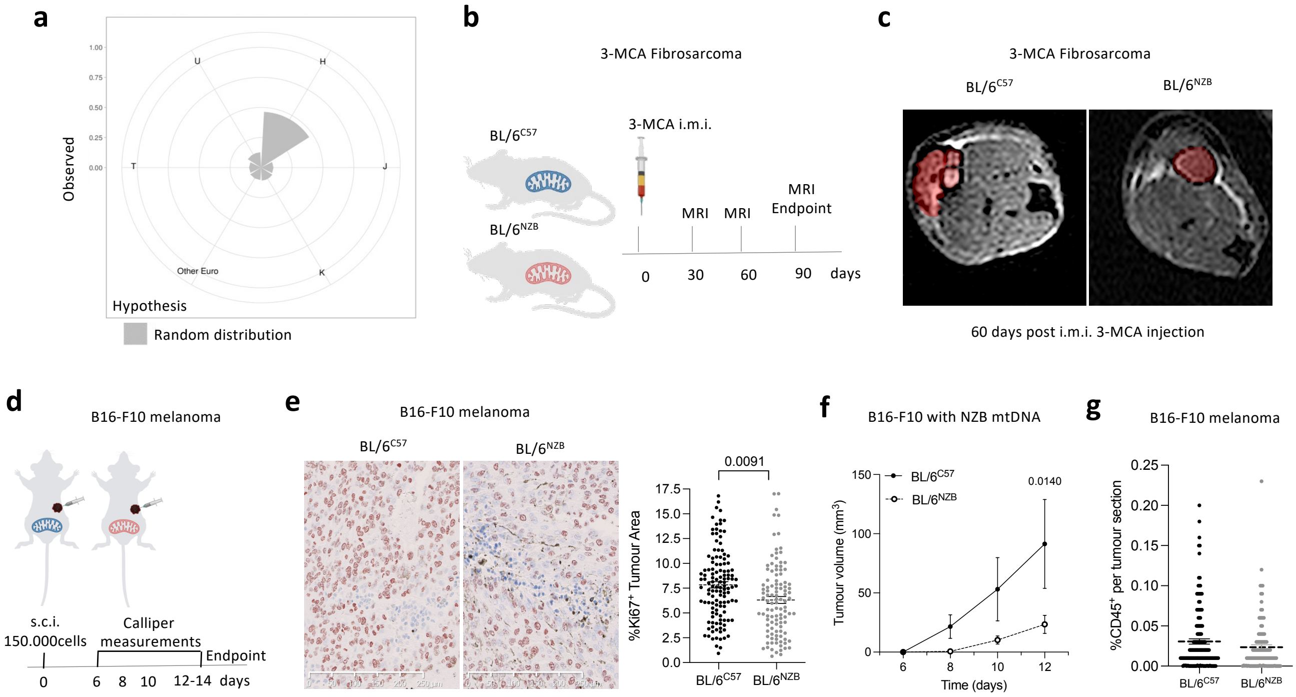


Extended Data Figure 1

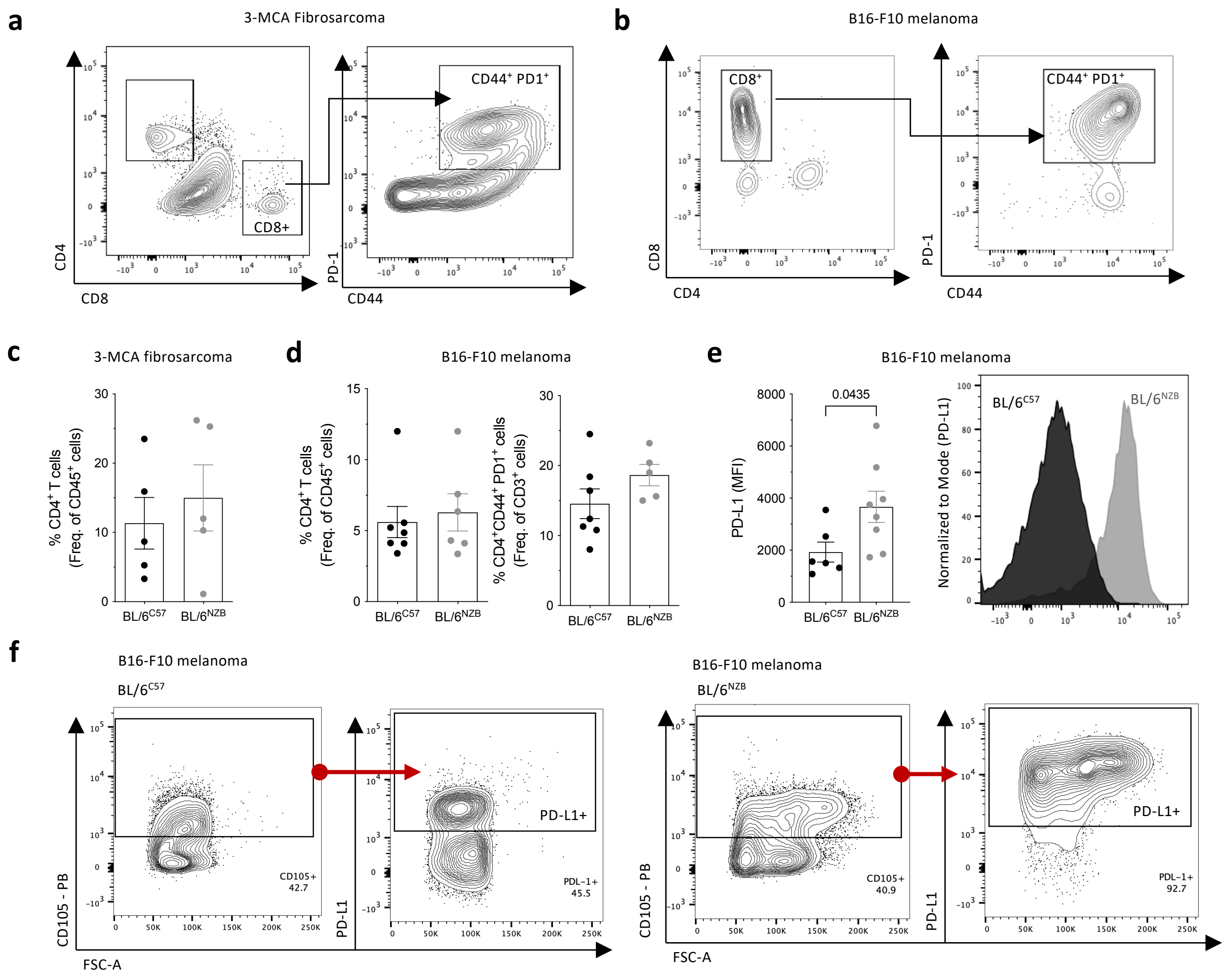


### Extended Data Figure 2

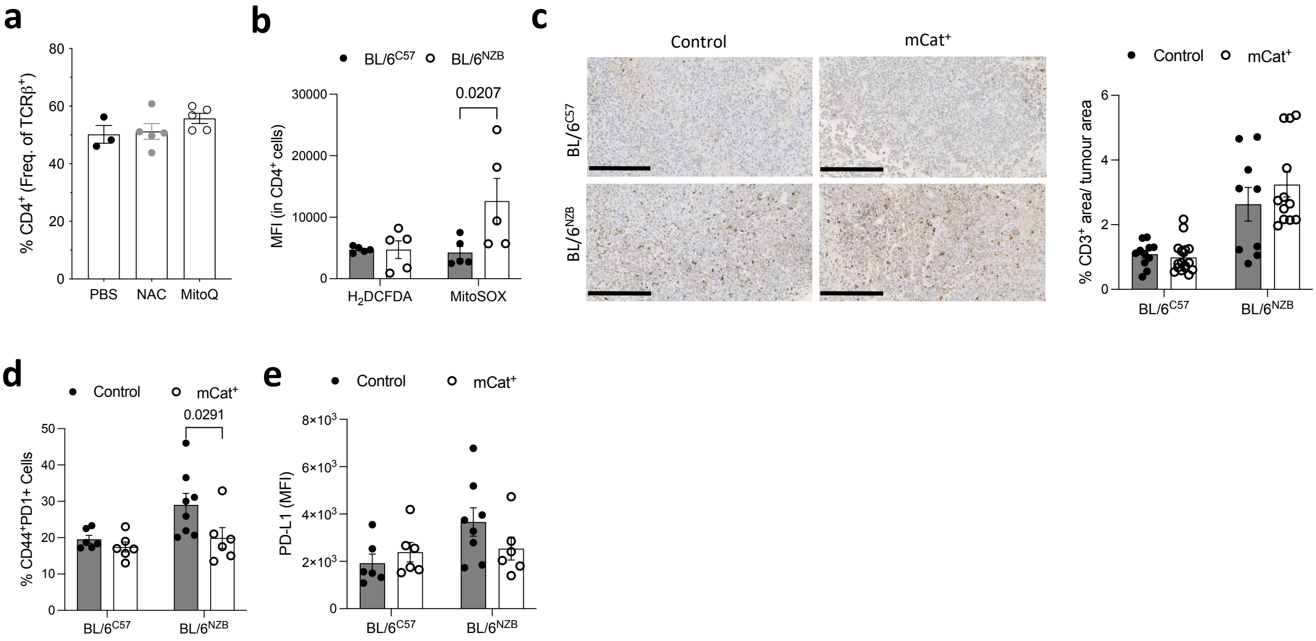


### Extended Data Figure 3

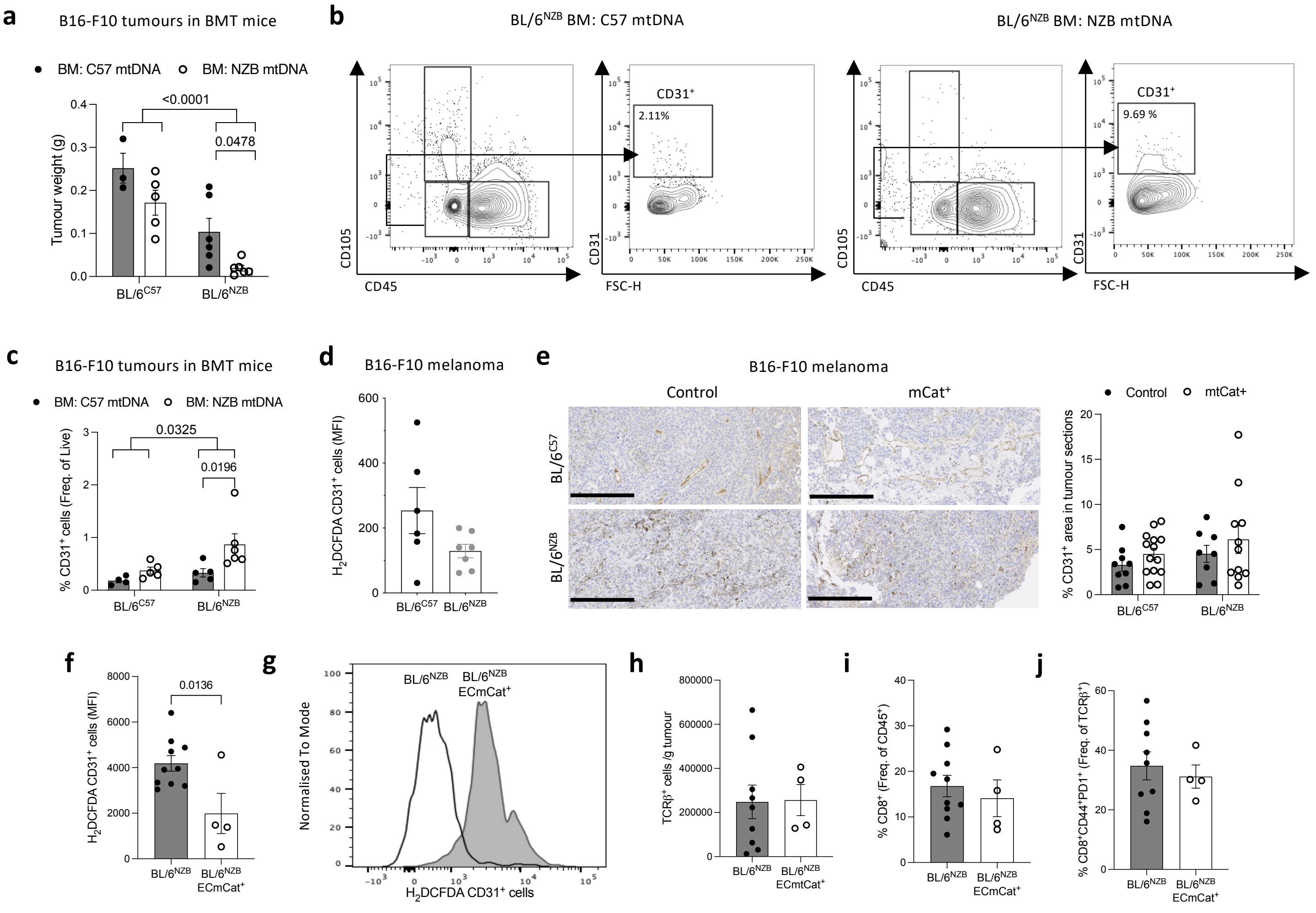


**Extended Data Figure 4**

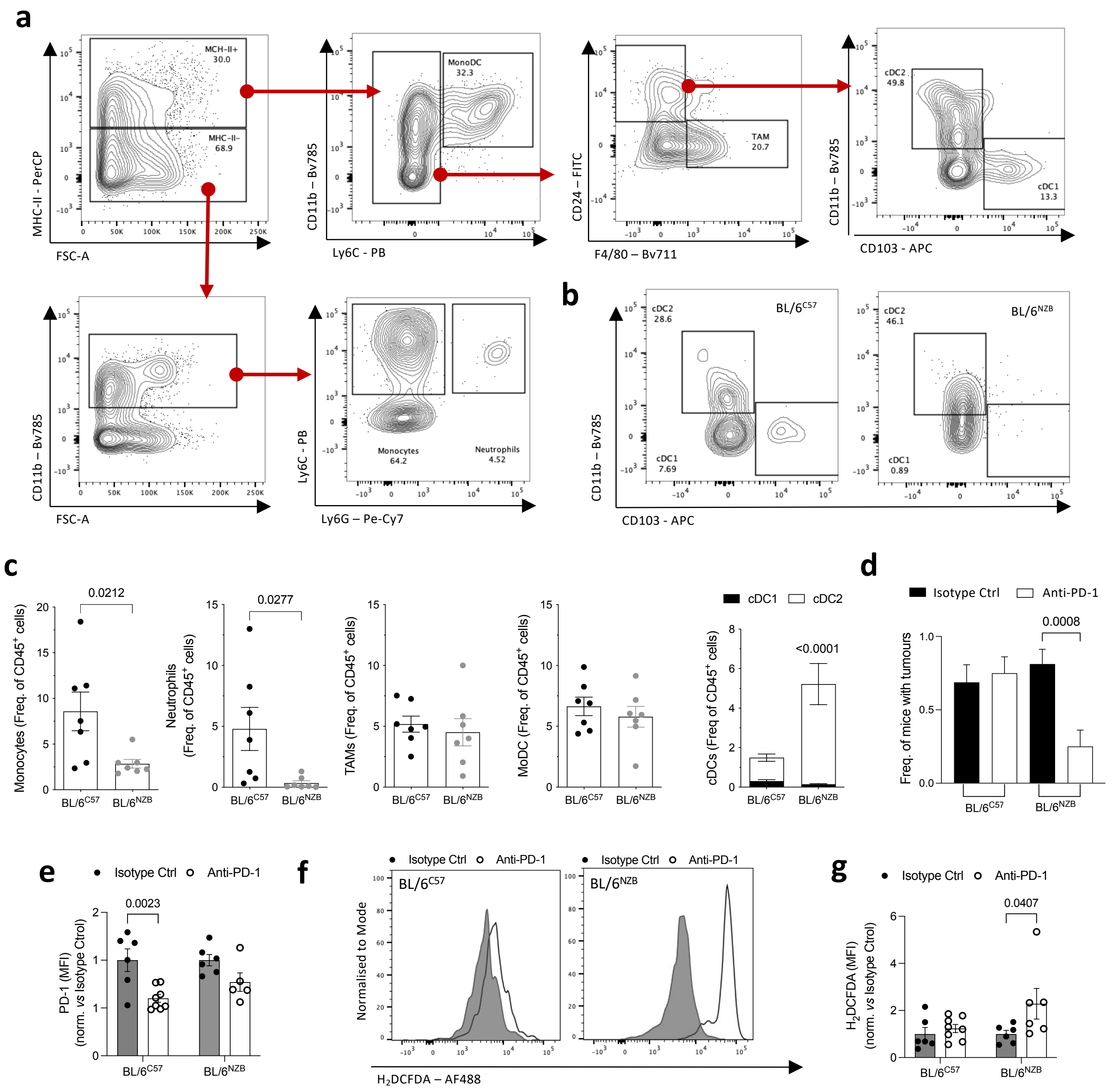


**Extended Data Figure 5**
