## Supplementary Table for "Mitochondrial genetics defines anti-tumour immunity through mitochondrial ROS and PD-1 signalling"

**Supplementary Table 1. Binomial test results for Macro-Haplogroup bias among Melanome patients.**

| HG | Hypothesis | Observed | Expected | p.value | adj.p* |
|---|---|---|---|---|---|
| **Other Euro** | Random distribution | 0.1037 | 0.1016 | 0.8131 | 0.813125 |
| **J** | Random distribution | 0.1004 | 0.1111 | 0.2555 | 0.511183 |
| **H** | Random distribution | 0.4637 | 0.4527 | 0.4386 | 0.657978 |
| **K** | Random distribution | 0.1086 | 0.0892 | 0.0115 | 0.068825 |
| **T** | Random distribution | 0.0963 | 0.1001 | 0.7033 | 0.813125 |
| **U** | Random distribution | 0.1273 | 0.1452 | 0.0395 | 0.118616 |

*False discovery rate corrected by Benjamini-Hochberg method.
